# Tracking West Nile virus dynamics using viral loads from trapped mosquitoes

**DOI:** 10.1101/2025.07.02.662782

**Authors:** Punya Alahakoon, Ian Marchinton, R. Tobias Koch, Joseph R. Fauver, James A. Hay

## Abstract

Mosquito-borne virus transmission is increasing globally. Due to a lack of effective vaccines or targeted treatments, mitigation relies on reducing human exposure to infectious mosquitoes. Entomological surveillance provides estimates of human risk by quantifying the prevalence of arboviruses in mosquito populations. Testing of pooled mosquitoes via RT-qPCR is often used to infer infection prevalence, however these data are often binarised and ignore semi-quantitative cycle threshold (Ct values) that may provide valuable information on virus kinetics, mosquito infectiousness, and ultimately human risk.

West Nile virus (WNV) is a globally distributed arbovirus and endemic to the United States. In this study, we analysed pooled Ct values from retrospective mosquito surveillance and found substantial variation that could not be explained by laboratory factors alone. To understand how underlying viral load and epidemiological dynamics might explain this variation, we developed a multi-scale model linking pooled Ct values to heterogeneous within-host viral kinetics in mosquitoes and birds and seasonal transmission dynamics.

Our modelling suggests that Ct values from positive pools might reflect a mixture of potentially infectious and non-infectious mosquitoes, indicating that a substantial proportion of PCR-positive mosquitoes might not contribute to transmission. We then developed a method to estimate mosquito infection prevalence using only pooled Ct values that distinguishes infectious from non-infectious prevalence. Estimates of overall prevalence were comparable to standard methods using binarised data and remained accurate at higher prevalences where standard approaches fail.

Overall, these findings demonstrate that entomological viral load data reflect biologically meaningful variation in mosquito infection status that is lost through binarisation. Accounting for heterogeneous mosquito viral kinetics can improve interpretation of surveillance data and estimates of transmission risk, with implications for arbovirus surveillance beyond WNV.

## 1 Introduction

West Nile disease is a vector-borne disease caused by West Nile virus (WNV) infection [1]. It is maintained in an enzootic cycle between mosquitoes and birds [1, 2], with humans as incidental hosts [3]. Since WNV was first isolated in 1937 in Uganda [4], the pathogen has circulated globally, including southern Europe, Africa, Asia, the Middle East and Australia [2]. Due to climate change and human-induced activities such as land-use, travel, and urbanisation, there have also been changes in the global range of mosquito species for vector-borne diseases, including WNV [5, 6, 7, 8]. Before its introduction to the United States (US) in 1999, the severity of the disease among humans was considered low or moderate [2]. Since its introduction, there have been more than 60,000 clinical cases of WNV disease reported in the US, more than 30,000 of which resulted in neuroinvasive disease, and nearly 3,000 deaths [9]. Human vaccines and treatments are not yet available. [10, 11]. Therefore, the mitigation of disease depends on reducing human exposure to infectious mosquitoes through vector control and public health messaging [11, 12, 13]. These disease-mitigation interventions must be informed by immunological, epidemiological, and environmental parameter estimates of human risk.

Arboviral surveillance comprises both epidemiological and environmental approaches [14, 15]. While epidemiological surveillance is essential to quantify the human disease burden, it is not generally predictive of outbreaks, as human cases are primarily subclinical, occur during peak transmission periods, and experience lags between diagnosis and exposure. Environmental surveillance is therefore a crucial early indicator of human risk [14]. This includes surveillance in birds and monitoring weather and climatic factors, though mosquito surveillance remains the most reliable leading indicator of human risk, as enzootic virus transmission is detected in mosquito populations weeks earlier than human cases are identified [16]. Vector-based WNV surveillance involves collecting vector species mosquitoes at regular intervals (e.g., weekly, monthly) and using molecular testing for the presence of WNV [14]. Data gathered through this process informs indicators of quantifiable risk such as vector abundance, infection rate, and vector index for implementing vector control efforts [14]. We focus here on vector-based surveillance and explore strategies to enhance its effectiveness in quantifying WNV prevalence in mosquito populations.

Mosquito-based surveillance detects WNV RNA in pools of trapped mosquitoes using reverse transcription quantitative polymerase chain reaction (RT-qPCR). RT-qPCR measures the amount of viral material in the pool as cycle threshold (Ct) values, which are inversely correlated with the quantity of viral RNA, though pool status is typically treated as positive or negative resulting in statistical information loss [17, 18]. However, it has been shown that using Ct values from pooled test data can improve prevalence estimation for SARS-CoV-2 infection, particularly in resource-constrained settings [19]. In addition, Ct values from virologic RT-qPCR testing vary over the course of an epidemic, reflecting the interaction of epidemic dynamics and within-host kinetics [20]. In the case of acute respiratory virus infections, this arises because infections are more recent on average during a growing epidemic, which coincides with typically higher viral load, whereas infections are less recent during a declining epidemic, coinciding with typically lower viral load. Thus, for SARS-CoV-2, the distribution of Ct values can be used to infer epidemic dynamics with some statistical advantages over treating tests as positive or negative [20].

WNV dynamics are well-characterized in mosquitoes through laboratory experiments [21, 22, 23]. These studies demonstrate that WNV titers in mosquitoes grow steadily following ingestion of an infectious bloodmeal, midgut infection, and viral dissemination. Importantly, mosquitoes are unable to clear viral infection and therefore maintain high viral titers throughout their lives [24, 25]. The substantial variation in viral quantities observed during cross-sectional entomological surveillance suggests more complex vector/virus interactions, and the precedence set by population SARS-CoV-2 testing in humans suggests that Ct value data for WNV in mosquitoes could inform metrics of disease risk to humans. However, there are substantial differences in the epidemiology and biology of WNV infection in mosquitoes with SARS-CoV-2 in humans. How the interaction between epidemic dynamics, within-host kinetics and sampling strategy affects the distribution of observed Ct values from entomological surveillance data for arboviruses such as WNV has not yet been explored.

Here, we analysed WNV mosquito surveillance data from Nebraska and Colorado, two high-incidence states with robust mosquito surveillance programs in the US. Weekly entomological data consisted of mosquito abundance, binary (positive/negative) results, and Ct values from mosquito pools. We sought to propose biological mechanisms which might explain observed variation in pooled Ct values. In particular, we observed a bimodality in the observed Ct value distributions, where the majority of the Ct values were low (high viral loads), but high Ct values (low viral loads) were consistently observed, which are unlikely to be explained by dilution or laboratory effects. To understand mechanisms which might explain observed variation in WNV Ct values over time, we developed a multi-scale agent-based model that integrated population-level dynamics and viral load kinetics of mosquitoes and birds. We implemented a synthetic sampling strategy to mimic mosquito surveillance and the RT-qPCR testing process, generating pooled Ct values consistent with surveillance data from Nebraska and Colorado. Based on this mechanistic model, we investigated if pooled Ct values can serve as a reliable data source for accurately estimating mosquito prevalence in a population.

## 2 Results

### 2.1 Substantial variation in WNV quantities from entomological surveillance data

We analysed entomological trap data tested for West Nile virus (WNV) using RT-qPCR [26] collected through routine environmental surveillance in Nebraska (statewide) from weeks 23-37 in 2022 to 2024, and Colorado (Larimer County) from weeks 23-37 in 2022 and 2023. In Nebraska, pools either combined *Culex pipiens/restuans/salinarius* or *Culex tarsalis*, whereas in Colorado, pools contained either only *Culex tarsalis* or *Culex pipiens*. For the purpose of measuring WNV prevalence, we combined data from all pools regardless of species. Ultimately, our data includes Ct values from thousands of pools of mosquitoes over 2 trapping seasons in Colorado, and 3 trapping seasons in Nebraska (Figure 1) See Supplementary Material S1 for a visualisation and distribution of Ct values of other species and to potential relationships between Ct values and other factors.

**Figure 1:**
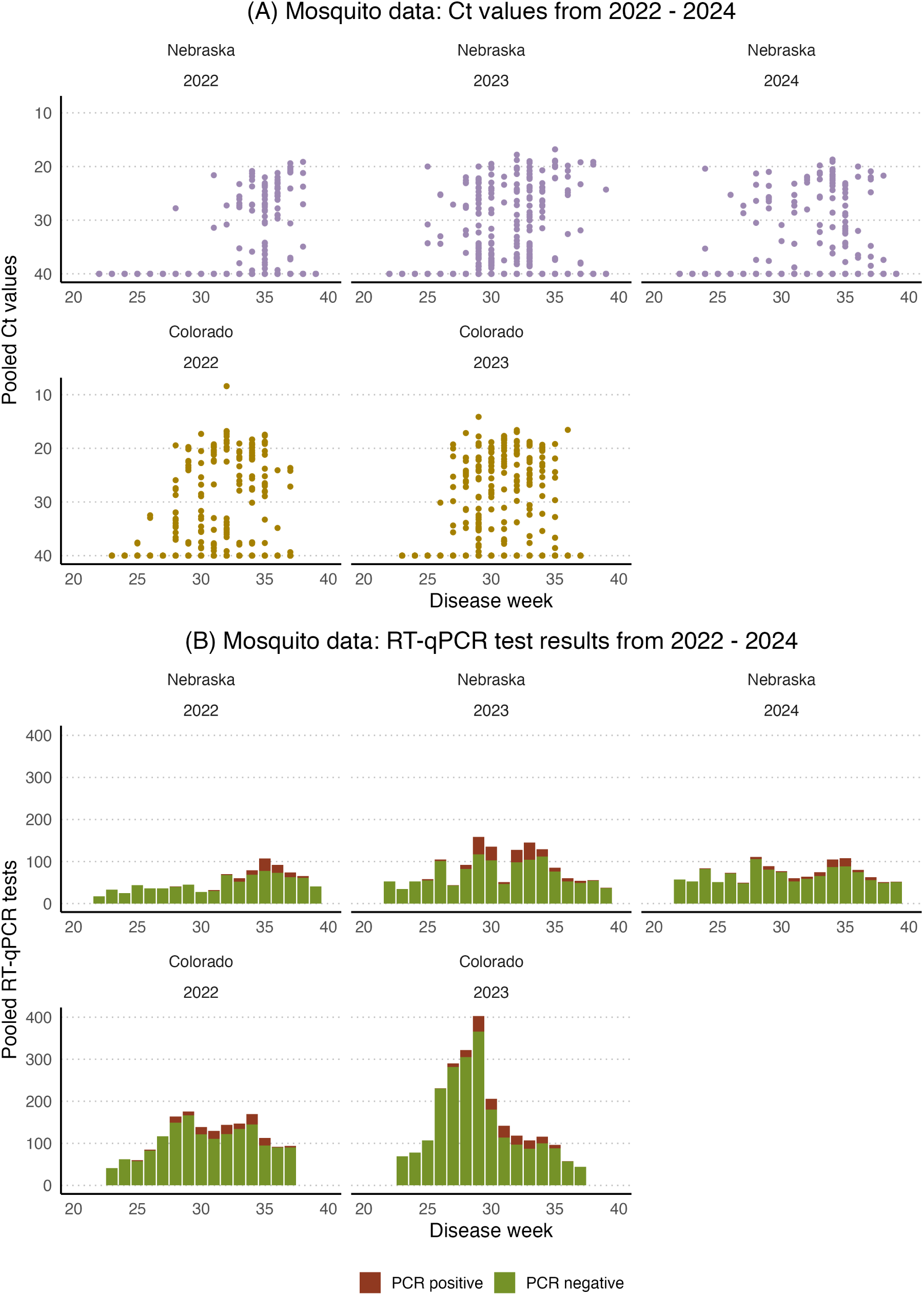
Observed data: **(A) :** Calculated pooled Ct value vs. disease week from 2022 to 2024. Ct values are presented from pools of mosquitoes separated by species, either *Culex pipiens/restuans/salinarius* or *Culex tarsalis*. **(B)** : RT-qPCR test results vs. disease week from 2022 to 2024.

We observed significant variability in Ct values within the limit of detection (Figure 1). Cross-validation of Ct values between laboratories and observation of dramatically different Ct values across all pool sizes, localities, and through time suggest that this variation is likely genuine as opposed to a laboratory artefact. First, the variation we observe is consistent across five trapping seasons and two states (Figures S5 and S6). The surveillance data presented in this study come from two independent laboratories, one of which is a certified state public health laboratory. Both laboratories implement rigorous controls, including no template controls in both extractions and RT-qPCRs.

Second, the variation in Ct values is dramatic, ranging from 10’s to 10’s of millions of copies of WNV RNA in individual pools (Figure S2). Although pooling samples would not reduce the total quantity of target nucleic acid in a sample, we tested whether our Ct value data may be biased by the pooling of mosquitoes leading to reduced sensitivity of the RT-qPCR assay. We hypothesised that if pooling had a strong effect on WNV Ct values, we would systematically observe higher Ct values and more false negatives in pools with more mosquitoes as the total amount of WNV RNA from a positive mosquito body would be diluted with additional WNV-negative mosquito tissue. Although a dilution effect was expected and explicitly incorporated into the model (Supplementary Material S4), we observed that the variation in Ct values remained consistent across pool sizes and weeks (Figure S2). A pool size of 50 is the most common pool size in our dataset and the Ct values from the Nebraska data ranged from 19.4-39.2 in pools of 50 mosquitoes, and 16.8-39.4 in pools of 25 or fewer mosquitoes (Figure S2). These results indicate that pooling mosquitoes is unlikely to reduce the sensitivity of RT-qPCR assays to detect WNV RNA.

Finally, to determine the reproducibility of these data, a subset of 143 WNV-positive pools from Nebraska were retested in a separate research laboratory at University of Nebraska Medical Center (UNMC). The average Ct value difference between samples tested in both labs was −2.26, or a *≈*10-fold difference in measured RNA quantity, with Ct values consistently being lower in the independent laboratory (Figure S3). A total of 14 samples tested negative at the UNMC laboratory, and the average Ct value of these samples was 38.24. Ct values from the UNMC laboratory were consistently lower, however the reduction was consistent across the entire range of Ct values. This is likely due to the original Ct values from Nebraska being produced using a multiplex assay that includes primers/probes for St. Louis encephalitis virus and western equine encephalitis virus as well as WNV, reducing the sensitivity of the assay for any one target [27]. The consistency of Ct values produced from both labs highlights the reproducibility of WNV Ct values from mosquito pools. Taken together, the Ct values produced from routine WNV surveillance represent genuine variability in WNV RNA from mosquitoes.

After ruling out laboratory effects, we were interested in investigating potential mechanisms which might explain observed variation in pooled Ct values. Viral loads in infected mosquitoes are driven by multiple factors, including infectious dose, temperature, virus genotype, and time point of collection during the extrinsic incubation period (EIP), or the time from virus ingestion to infectiousness in mosquitoes. [28, 29, 30, 23]. However, we expect host-seeking mosquitoes infected with WNV to have high viral loads (low Ct values) due to viral replication during the gonotrophic cycle prior to host seeking for a subsequent blood meal. Given the substantial variability observed in Ct values from host-seeking mosquitoes (Figure 1), we hypothesised that this variation may be due to biological and epidemiological factors, such as variability in the viral kinetics in mosquitoes and mosquitoes being captured at different times post-infection. Accordingly, we developed an agent-based model (ABM) that captured these mechanisms, enabling us to compare predictions consistent with these hypotheses to the observed data.

We used the ABM to simulate population-level and viral load dynamics of mosquitoes and birds (Figure 2; see Section 5 and Supplementary Material S2-S5 for further detail). Overall transmission dynamics were based on the Ross-Macdonald model[31, 32], describing the transmission cycle between bird hosts and mosquito vectors (Figure 2 (A), (B) & (D). We embedded within-host models of virus kinetics for mosquitoes and birds describing how WNV viral loads change over time since infection and vary between individuals (Figure 2 (C)). The model incorporates host seeking behaviour and key environmental factors such as seasonality and overwintering. Using this model, we simulated the trapping and pooling process to generate Ct values from pools of up to 50 mosquitoes to match the real-world surveillance data in Nebraska (Figure 2 (E)). The outputs of the model are simulated pooled mosquito Ct values for WNV, collected weekly over the course of multiple seasons and reflecting underlying WNV infection prevalence in the mosquito population (Figure 2 (F)). We parameterised our agent-based model to recreate WNV prevalence patterns observed in the Colorado and Nebraska surveillance data (Supplementary Material S2-S5).

**Figure 2:**
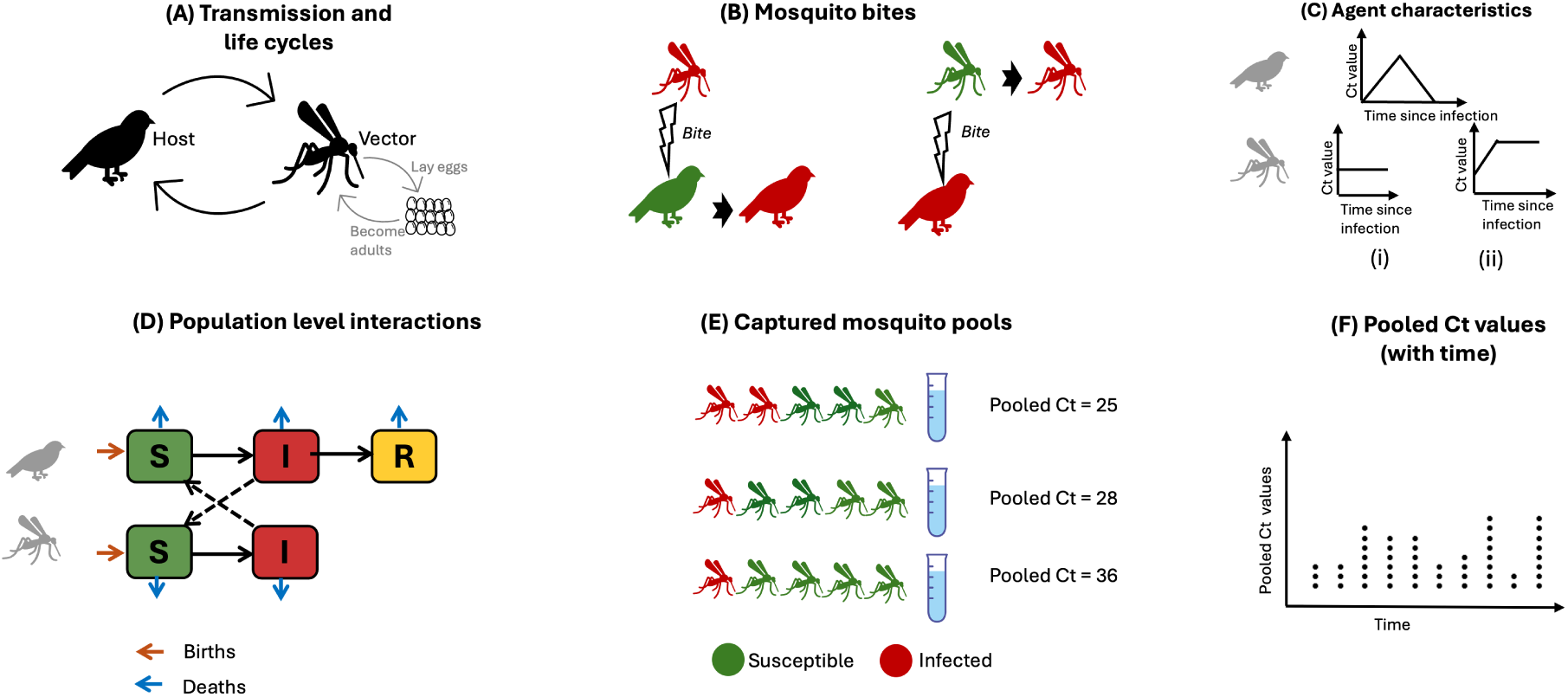
Overall structure of the agent-based model (parameters were derived from the literature; see Supplementary Material S2, S3 and S4). **(A) :** Transmission cycle of WNV between birds (hosts) and mosquitoes (vectors). Female mosquitoes lay eggs, which hatch after an average of 8 days, before becoming female adult mosquitoes that contribute to the transmission cycle. **(B) :** Transmission occurs with some probability when an infected female mosquito bites a susceptible bird, or a susceptible female mosquito bites an infected bird. **(C)** : Within-host viral kinetics model. Infected birds follow an acute viral load trajectory with rising and clearance phases, initiating from the RT-qPCR limit of detection. Female mosquitoes may follow one of two different viral kinetics models: the mosquito’s initial viral load is proportional to the infector bird’s viral load at the time of bite and either leads to a non-replicating viral trajectory (static viral load) (i) or a replication-competent infection (dynamic viral load) (ii). Note the Ct value scale is inverted, with lower Ct values depicting higher on the scale. **(D) :** Overall population-level dynamics of the agent-based model. If bitten by an infected mosquito, susceptible birds (*S*) can become infectious (*I*). Once infectious, they will eventually recover (*R*) with lifelong immunity following their infectious period. Susceptible mosquitoes (*S*), if they bite infectious birds, will become infectious (*I*) for the rest of their lifespan. Births and deaths are possible in all these states for both birds and mosquitoes. **(E) :** Simulation of routine mosquito trapping and pooled testing accounting for blood meal-seeking behaviour. For each pool, a combined pooled Ct value is calculated as the mean viral load of captured WNV-positive mosquitoes, diluted by the contribution of WNV-negative mosquitoes. **(F) :** Example population-level Ct dynamics from the simulation. Note the Ct value scale is inverted, with lower Ct values depicting higher on the scale.

We ran the simulation four times, and each simulation was run for twenty-six years, with the first year excluded as a burn-in period due to the atypical dynamics expected following initial WNV introduction into a fully susceptible population [33, 34]. This resulted in 100 one-year WNV epidemics (Figure 3). Full simulation dynamics are shown in Supplementary Material S6. The model generated peak WNV infection prevalence of 3-4% in mosquitoes and peak infection incidence of 0.17-0.36% in week 29-36, coinciding with the summer months in the US (Supplementary Material S6). Simulated pooled Ct values were calculated in weeks 22 to 39, matching the surveillance data from Nebraska (Figure 3).

**Figure 3:**
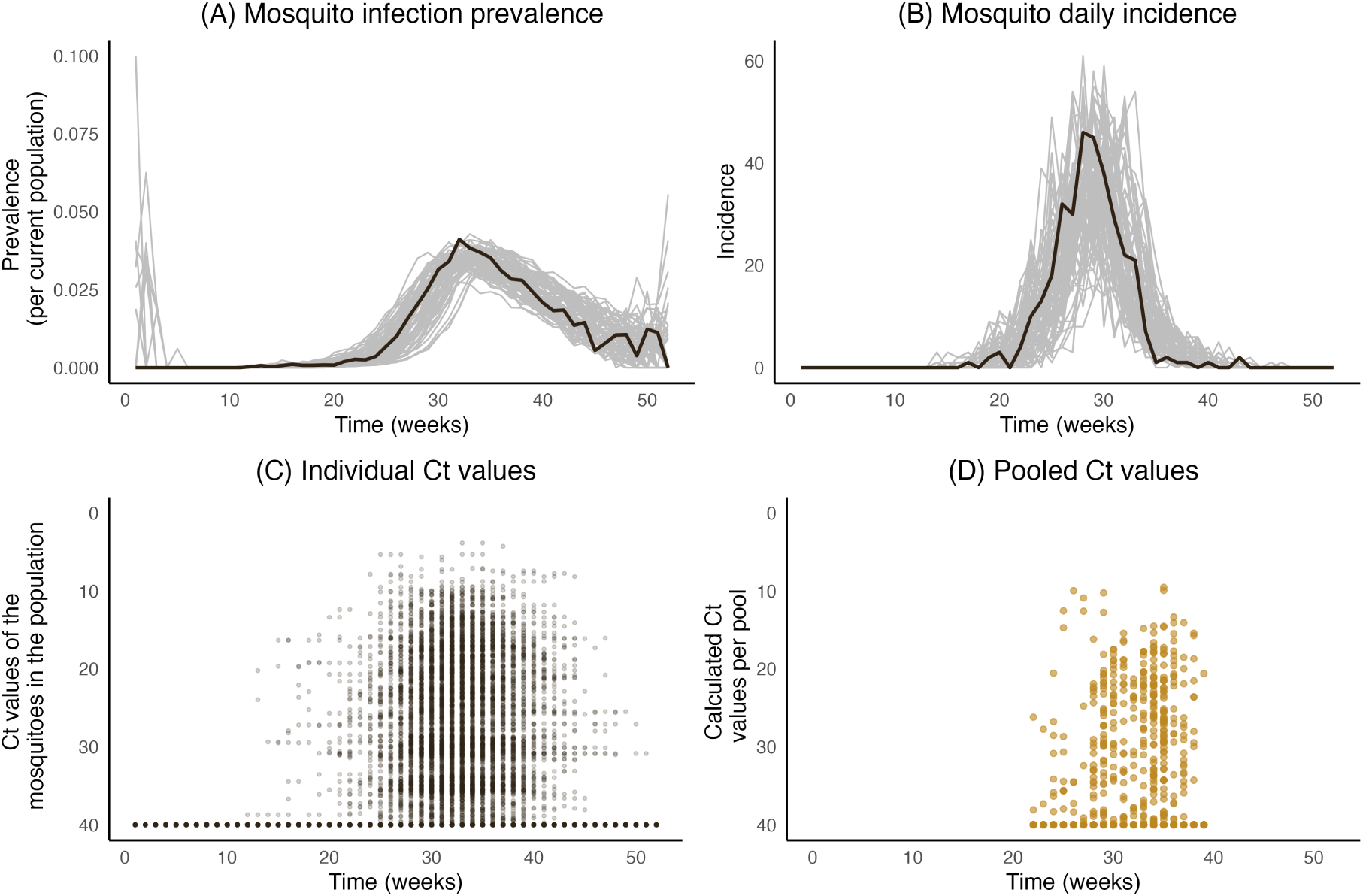
Simulated population-level WNV infection dynamics in mosquitoes. Each line represents one year of simulated data, and the bold black line highlights the simulated trajectory used for (C). **(A) :** Weekly mosquito infection prevalence per capita. **(B) :** Weekly mosquito infection incidence per capita. **(C) :** Individual-level Ct values of the female mosquito population from one year of a single simulation. Each point represents a Ct value from a single randomly selected mosquito, representing perfect observations of the entire population at all times. **(D) :** Calculated pooled Ct values from the mosquito population based on the individual-level Ct values in (C)

Unlike acute viral infections in humans, where viral load initially increases but then decreases as the infection is cleared [35, 36, 37], productively infected mosquitoes quickly reach and maintain a high WNV viral load once infected. This is reflected in the population Ct value distribution, which does not change over time in both individual-level and pooled testing (Figure 3 C & D). For time variation in the population-level mean Ct value to arise, two conditions must be met [20]. First, there must be variation in the time-since-infection distribution over time, which is guaranteed in an epidemic; the average time-since-infection is lower in a growing epidemic than a declining one. Second, the expected Ct value must vary over time-since-infection to allow recent infections to be distinguished from older infections. Although the first condition is met here, the variation in viral load over time-since-infection is small for WNV infection in mosquitoes, and thus the population-level Ct distribution is static. The mosquito pool Ct value data in this study come from specimens collected using CDC Light Traps that are baited with CO_2_, specifically targeting host-seeking mosquitoes. Vertical transmission is not factored into our model as it has been reported to occur infrequently [38], thus, for a WNV-positive mosquito to be captured in the pool, it must have already obtained one blood meal from an infected bird and be seeking its next blood meal, which introduces a delay between infection and being captured. As the infected mosquitoes progress from increasing viral load to a high set-point viral load, temporal trends in pooled Ct values primarily reflect time-varying infection prevalence and the number of infected mosquitoes in each pool. The remaining non-temporal variation in pooled Ct values reflect individual-level variation in mosquito set-point viral loads.

### 2.2 WNV population-level viral loads can be explained by lack of viral replication in mosquitoes

The model assumes that the mosquito’s initial viral load is proportional to the infector bird’s viral load (see Section 5.3 and Supplementary Material S3 for further details on the bird viral kinetics model) at the time of bite, and this may lead to a mosquito infection with either a static viral load reflecting non-replicating virus (non-productive infection), or a dynamic viral kinetics curve reflecting replicating virus (productive infection), until death or capture. High Ct value samples might arise from mosquitoes harbouring low levels of virus in their bodies without active replication occurring, for example, by ingesting non-replicating viral material from the blood of infected birds. This would have implications for interpreting positive pools, as it would suggest that not all positive pools reflect actively infected mosquitoes. Note that these are distinct from false positives which refer to PCR positive pools containing no WNV viral material. To test this, we systematically varied two model parameters: the proportion of viral load ingested and the probability of each mosquito’s post-infection trajectory being productive or non-productive. We compared the resulting distribution of simulated pooled Ct values to the observations in Nebraska and Colorado from 2022 to 2024 to identify parameter combinations most consistent with the real data.

Varying these two parameters changed the distribution of pooled Ct values substantially (Supplementary Material S7). We found that the simulated pooled Ct values aligned well with the observed data when the percentage viral load inherited from birds was 100% and the probability of a productive or non-productive infection in the mosquitoes was 0.5, capturing the bimodal distribution of low Ct values (from productively infected mosquitoes) and high Ct values (from non-productively infected mosquitoes) (Figure 4 (B), (C) & (D); Supplementary Material S7.1, Figure S14 for Ct distributions of viral inheritance probability vs. model change probability and Figure S22 for the accuracy and confidence-interval coverage across different productive infection proportions). Hereafter, our analysis is based on the assumption that 100% of the viral load is ingested from the bird to the mosquito at the time of a bite and that there is an equally likely chance that the post-infection trajectory of the mosquitoes is static or dynamic.

**Figure 4:**
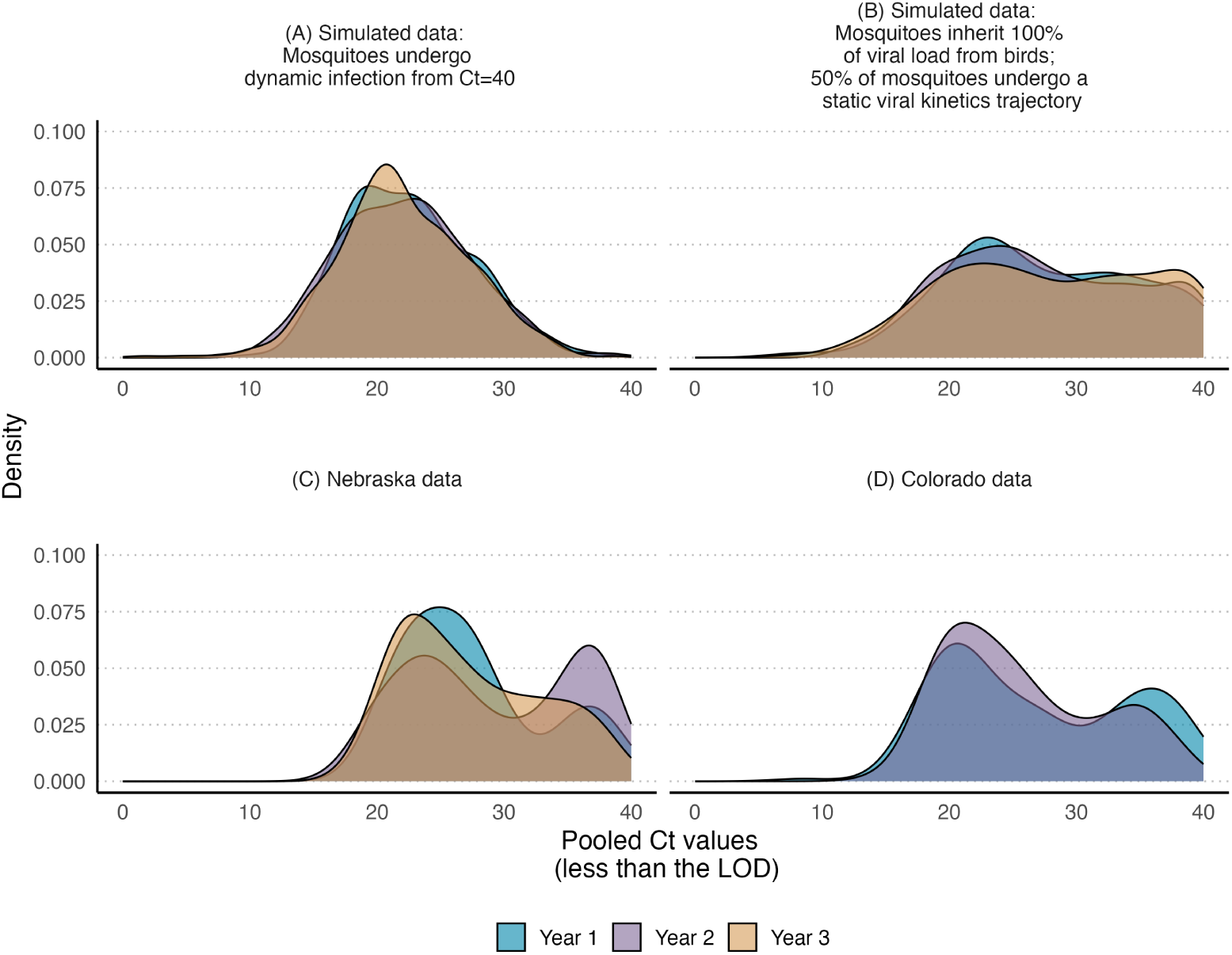
Distributions of the pooled Ct values within the limit of detection. **(A) :** simulated Ct values by assuming all mosquitoes undergo dynamic infection from Ct=40. **(B) :** simulated Ct values assuming mosquitoes inherit 100% of viral load from birds, and 50% of mosquitoes undergo a static viral kinetics trajectory. The bottom left and right panels. **(C) and (D):** the density of pooled Ct values from Nebraska and Colorado states over three years, respectively.

As a sensitivity analysis, we also examined whether post-infection viral-load decay in mosquitoes could plausibly explain the observed Ct distributions (Supplementary Material S7.2, Figure S15). Across all scenarios evaluated, introducing a non-negligible decay rate did not produce Ct distributions similar to those observed in Nebraska and Colorado. These results indicate that any substantial decline in viral load after infection is unlikely, and that assuming minimal or no decay provides a model that is most consistent with the empirical Ct data.

### 2.3 Calculating WNV infection prevalence using pooled Ct values

Next, we developed a method to estimate WNV infection prevalence in mosquitoes using pooled Ct values from captured mosquitoes. Based on a previous method developed for SARS-CoV-2 prevalence estimation, the method uses the expected distribution of Ct values for pools of a given size, a given number of positives, dilution effects, measurement error and variation in viral loads from infected mosquitoes [19]. We extended the method to treat overall infection prevalence as a mixture distribution of productive and non-productive infections, allowing us to estimate both the overall prevalence and proportion of infections which are productively infected. We fitted smoothed kernel density curves to our simulated viral loads to generate empirical distributions of Ct values for each combination of pool size, number of productively infected mosquitoes and number of non-productively infected mosquitoes. Our approach uses these density curves to construct a likelihood function for the population prevalence and proportion productively infected, conditional on the Ct values from multiple pooled test results. Using a maximum likelihood estimation method, we estimated the prevalence per week and the corresponding confidence intervals (more detail in Section 5.4 and Supplementary Material S8).

We assessed the accuracy of the Ct value-based method in recovering the true infection prevalence and proportion productively infected using the simulated pooled Ct values (Figure 5, Figures S16 and S17). We compared estimates of the overall WNV infection prevalence using our method (Figure 5 D) to estimates from the CDC *PooledInfRate* R package [39, 40], treating the pooled test results as positive or negative (Figure 5 E). We binarised the Ct values based on whether or not the values were below the limit of detection (Figure 5 B).

**Figure 5:**
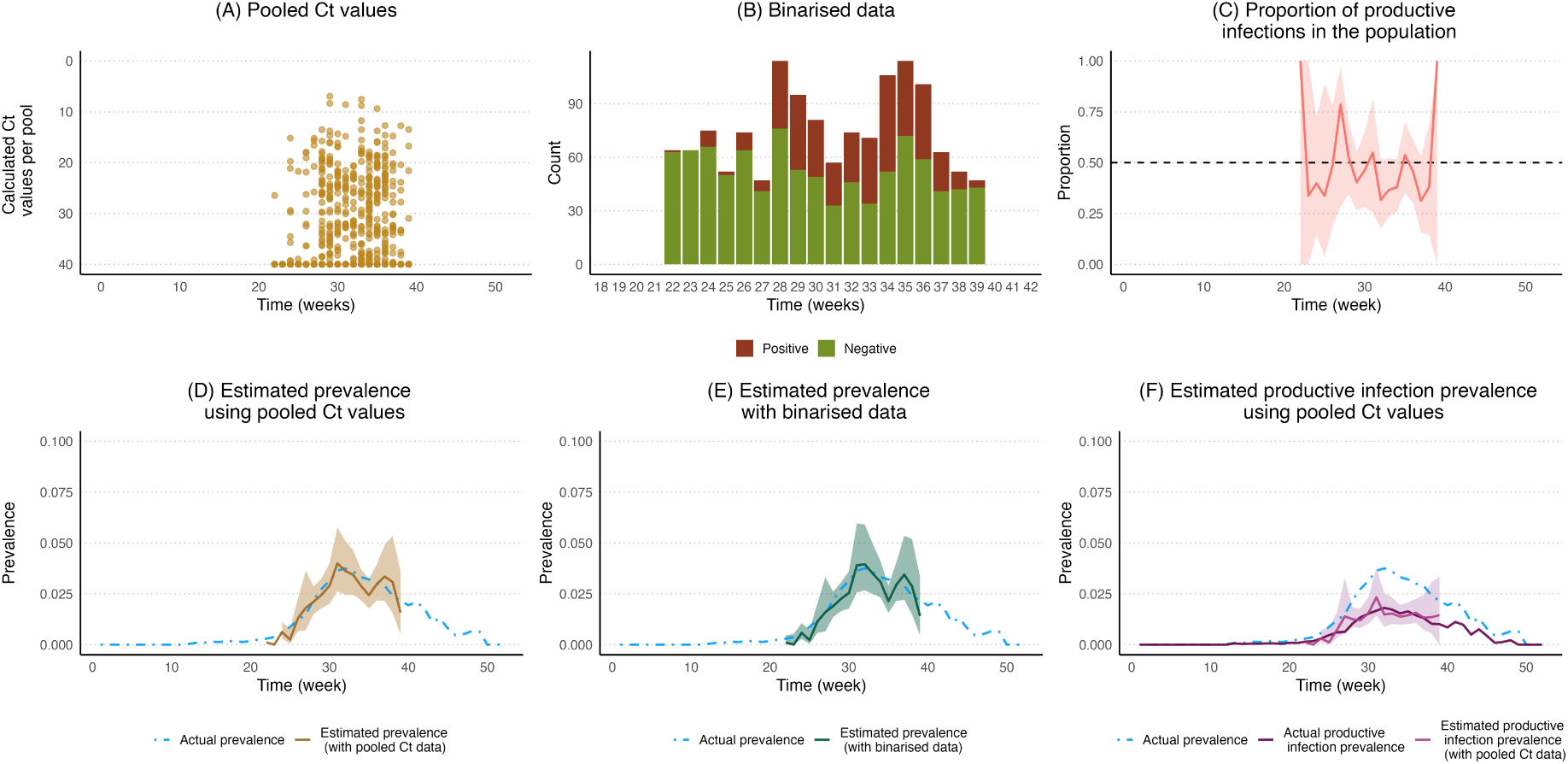
**(A)**: Weekly pooled Ct values from the captured mosquitoes (using the sampling procedure used in the agent-based model). Each dot represents the Ct value calculated per pool. **(B)** : Binarised data based on the pooled Ct values in (A). Binarisation was based on whether or not the pooled Ct values were within the limit of detection. **(C)** : Estimated proportion of productive infection using the pooled Ct values. The pink ribbon shows the 95% confidence interval. **(E)** : The estimated weekly WNV prevalence among mosquitoes using the binarised data with the *PooledInfRate* R package (green line). The green ribbon shows the 95% confidence interval. These values are compared with the true prevalence from the simulation (blue dashed line). **(F)** : Estimated productive infection prevalence using pooled Ct values. The purple ribbon shows the 95% confidence interval.

Overall, both estimation methods yielded similar results (see Figure S19 for prevalence estimation for multiple simulations) and were able to accurately estimate the mosquito prevalence when sufficient data were available. Point estimates were sometimes inaccurate under both estimation methods, likely due to sampling variation where the sample is a poor representation of the population prevalence. However, the 95% confidence intervals contained the true prevalence values. In the 10 simulation runs, each of a single year, included in this analysis, weekly coverage was first calculated within each run as the proportion of weeks for which the confidence interval contained the true prevalence, separately for the binary data-based and pooled Ct-based methods. Then the mean of these proportions were taken across the runs. These values were 0.917 [(0.790, 1.000) 95% quantiles] for the binary data-based method and 0.933 (0.846, 1.000) for the pooled Ct-based method. Similarly, we defined the weekly error as the difference between true and estimated prevalence, and then the mean error was calculated across all week-level estimates over all simulated years. The mean error was 0.0000816 (−0.0232, 0.0126) for the binary data-based method and −0.00117 (−0.0225, 0.0109) for the pooled Ct-based method. The pooled Ct-based method also gave accurate estimates for the proportion of productive infections and therefore of the prevalence of productive infections (Figure 5 C & F).

#### 2.3.1 Potential effects of pool size and number of pools on the accuracy of prevalence estimation

We next assessed the statistical power, coverage probability (whether the true prevalence was within the 95% confidence interval) and bias (difference between the MLE estimate and true prevalence) of the Ct-based and binary-based methods by simulating multiple datasets with different pool and sample size combinations, assuming true prevalence values ranging from 0.001 to 0.8 (Supplementary Material S9). Both methods generally performed well (Figure 6), though the *PooledInfRate* package could not estimate the prevalence effectively as the true prevalence and pool size increased. For example, with 50 mosquitoes per pool, the *PooledInfRate* failed to estimate the prevalence beyond 0.155. In contrast, using pooled Ct values allowed prevalence to be estimated consistently, regardless of the actual prevalence in the mosquito population. This is because with large pool sizes, almost all pools include at least one infected mosquito and are thus positive, whereas the Ct value-based method is still able to distinguish between higher and lower prevalences even if all pools are positive. The Ct-based method was unbiased at all pool sizes and number of pools, and became more accurate both with increasing number of pools (up to 500 pools) and with larger pools of up to 200 mosquitoes (Figure S20). Furthermore, we found that coverage was very high for both the Ct-based and binary-based estimates for smaller pools, though coverage of the binary-based estimates degraded as pool size increased, particularly with a large number of pools (Figure S21) when the true prevalence increased.

**Figure 6:**
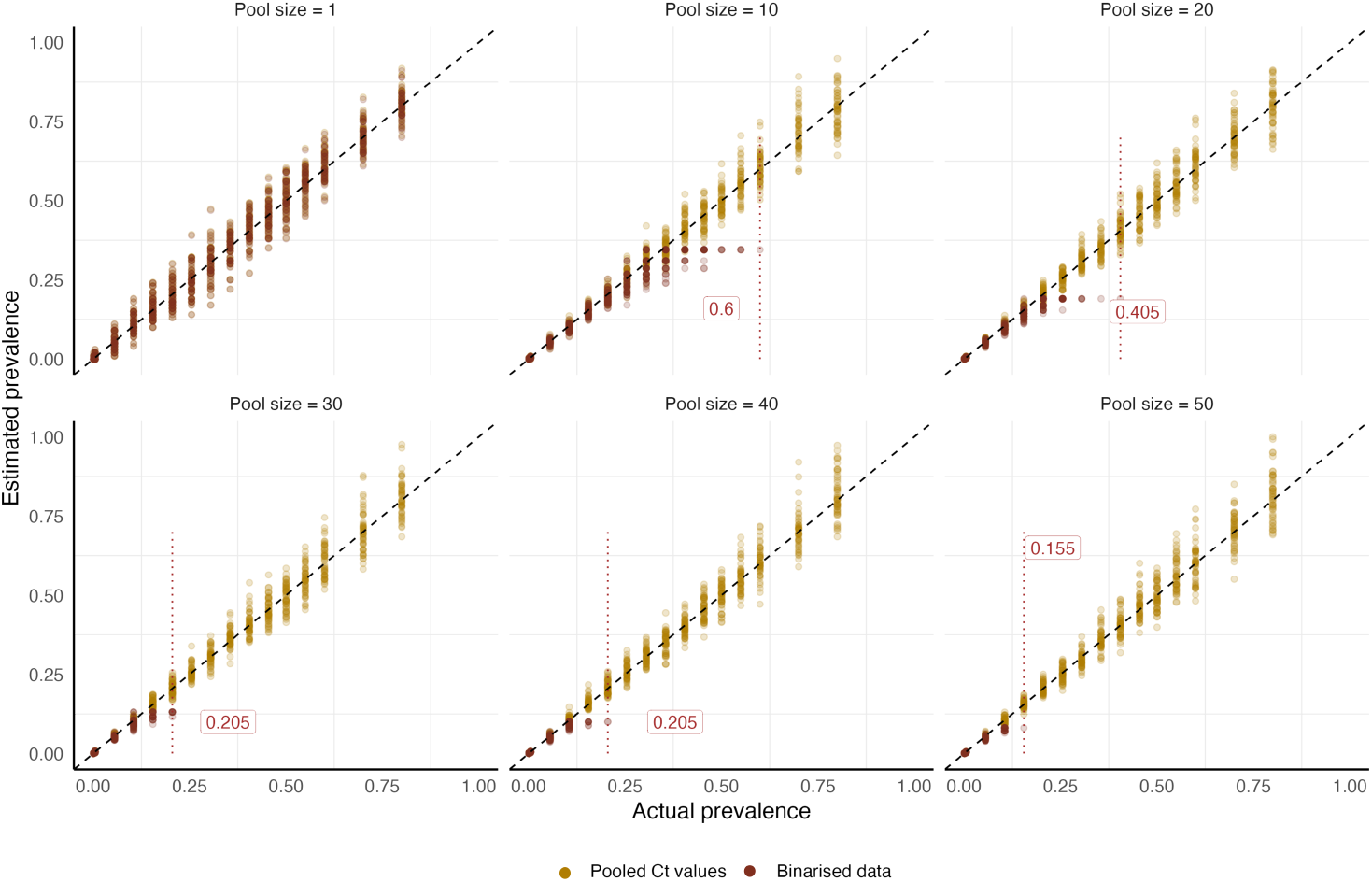
Comparison of prevalence estimates based on the Pooled Ct values and the *PooledInfRate* R package with varying pool sizes. The beige colour dots present maximum likelihood prevalence estimates calculated from pooled Ct values from a different simulation. The estimated prevalence calculated using the binarised data is in dark red. Each panel presents the estimates based on 100 pools containing up to 50 mosquitoes (pool size). The black dashed line represents perfect estimates. Labels along with the red dotted lines illustrate the maximum prevalence level at which the *PooledInfRate* R package returns accurate estimates.

## 3 Estimating WNV prevalence in the mosquito population based on RT-qPCR testing data from Nebraska and Colorado

Using the simulated pooled Ct values distributions from the model (Supplementary Material S8), we estimated the mosquito infection prevalence for Colorado and Nebraska from 2022 to 2024, comparing our Ct-based method to estimates based on binary pooled status (Figure 7). Across all study years, the binary and Ct-based methods produced broadly similar prevalence estimates. Using the Ct-based approach, the peak WNV prevalence was estimated at 0.015 [(0.009, 0.022) 95% confidence interval] in Nebraska in 2022 and 0.012 (0.007, 0.018) in Colorado in 2023. We observed that the widths of the confidence intervals were generally similar for both methods, though the Ct-based method showed greater precision when the underlying estimated prevalence was zero.

**Figure 7:**
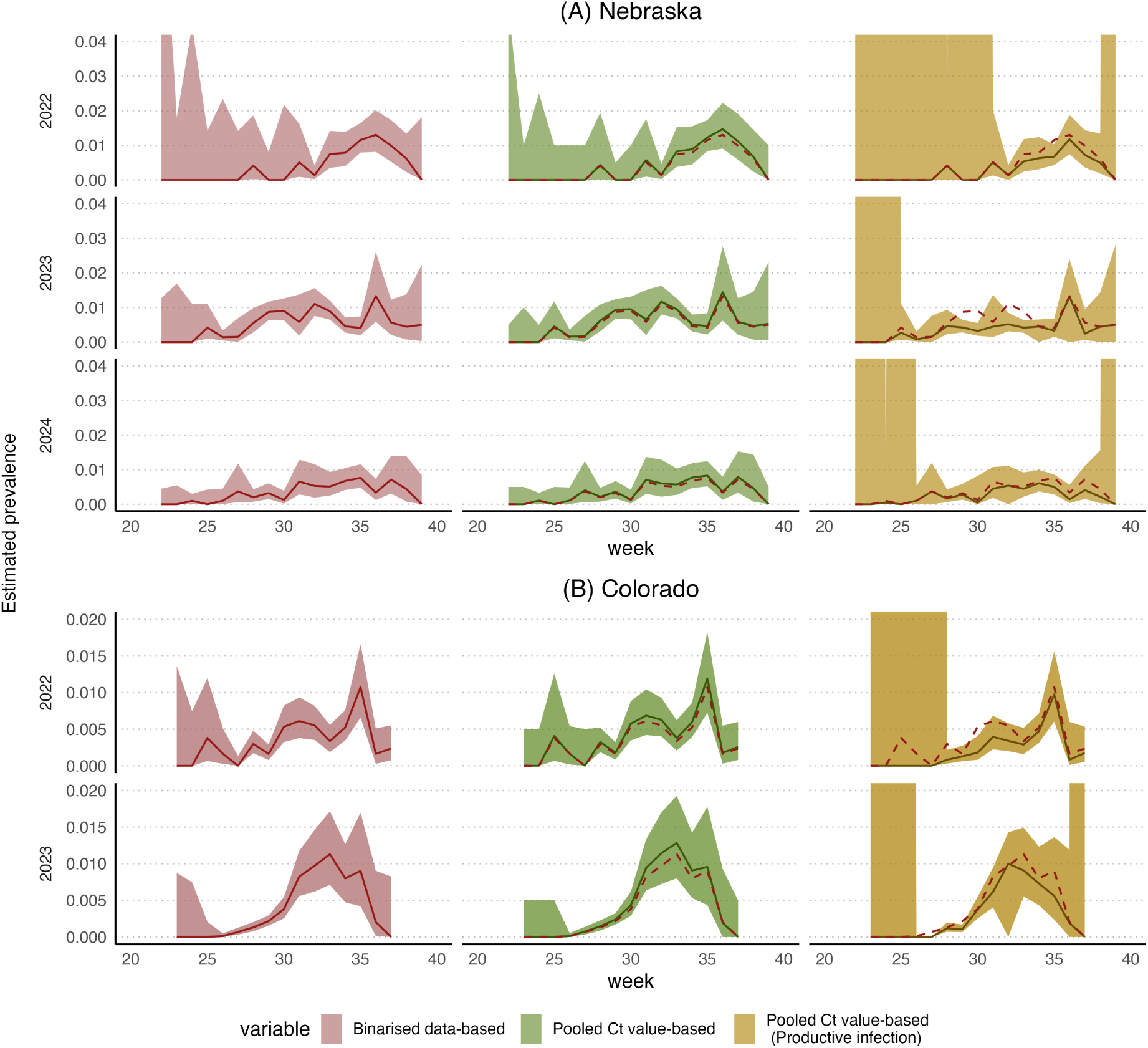
Overall prevalence estimation per year for Nebraska and Colorado from 2022-2024 using pooled Ct values (green) and binarised data (red), and the estimated productive infection prevalence (beige). The lines show mean estimates and the ribbons are 95% confidence intervals. Red dashed lines are the estimated prevalence based on binarised data.

A benefit of our Ct-based method is that we are able to estimate the productive (viral growth) and non-productive (no viral growth) mosquito infections separately (Figure 7). We therefore estimated the productive infection prevalence for each week for Nebraska and Colorado and compared it to the overall infection prevalence (Figure 7). We estimated the prevalence of productive infections to be around 50% of the overall infection prevalence across all weeks, but with high variability, likely due to uncertainty in the proportion parameter when overall infection prevalence was low and few pools were available (Figure S24), However, in some weeks, we estimated that most infections were productive. For example, 2023 week 39 (prevalence estimate 0.005 [(0, 0.028) 95% confidence interval]) for Nebraska and 2023 week 30 (prevalence estimate 0.004 (0.003, 0.005)) for Colorado had a maximum estimated productive infection proportion of 1 (Tables S2 and S3). These results are based on the model structure and the underlying assumptions we used and they may be affected by model misspecification, including incorrect assumptions (see [41] for example).

## 4 Discussion

In this study, we used WNV Ct values generated from routine entomological surveillance to model complex vector/virus interactions and improve estimates of WNV prevalence in mosquito populations.

We developed an agent-based model capturing transmission dynamics between mosquito vectors and bird hosts, heterogenous individual-level viral kinetics, and a realistic model for trapping and pooling mosquitoes. Based on comparison between our simulated data with real-world surveillance data from Nebraska and Colorado, we suggest that variation in WNV Ct values from mosquito pools might be explained by detecting a mixture of productively infected mosquitoes with high viral load and mosquitoes harbouring low-levels of non-productive viral material. Furthermore, we developed a new method for estimating WNV infection prevalence, distinguishing between the overall prevalence of productive versus non-productive infections, using Ct values from mosquito pools.

Data from entomological surveillance over five seasons in Nebraska and Colorado showed substantial variation in Ct values from mosquito pools tested positive for WNV RNA. We determined that laboratory artefacts or substantial dilution effects due to pooling were unlikely to account for the variation observed. Unlike SARS-CoV-2 infection in humans, where individuals experience a viral growth phase before generally clearing the virus [42], WNV viral loads in infected mosquitoes rapidly reach and remain at high levels following infection [1, 22, 43, 44], and thus the population-level Ct value distribution is not correlated with the epidemic growth rate. Sensitivity analyses assuming viral load decay following infection in mosquitoes were not compatible with the observed data. Nonetheless, we showed that population-level viral loads still reflect the interaction of epidemic dynamics and within-host viral kinetics; binarising pooled virologic surveillance data masks this information, which may be important for correct interpretation.

Multi-scale modelling techniques [45] allow individual-level biology and variation to be propagated into population-level dynamics. WNV infection in mosquitoes is a complex process where, upon ingestion of an infectious bloodmeal, the virus must initiate an infection in the epithelial cells of the mosquito midgut, replicate, and disseminate through the haemocoel prior to infecting salivary glands to perpetuate transmission [46]. Laboratory studies show WNV infection and growth rate in mosquitoes are not uniform, and the ability of mosquitoes to transmit virus is influenced by multiple factors [28], including temperature[29], host viremia [47], and genotype of the virus [23] and vector [22]. These factors result in heterogeneous mosquito infection kinetics, which is captured by our agent-based model.

Our results suggest that some low-viral load pools may contain infected mosquitoes that are incapable of transmission. This could be explained by a variety of factors, including mosquitoes being refractory to infection due to tissue escape barriers [48, 49, 50], heterogeneity in viral dynamics of infected birds [51], or timing of mosquito collection in relation to the EIP. The latter is unlikely to account for the observed variation, as all positive samples come from host-seeking mosquitoes collected via CDC Light Traps, meaning that WNV-positive mosquitoes were exposed to viraemic blood meals days or weeks prior to their collection. The low viral amounts detected in mosquito pools may be the result of the virus failing to initiate a productive infection, potentially due to virus and vector factors, implying that mosquitoes with high Ct values may not be able to transmit WNV. Our results highlight the importance of considering variation in the individual-level virus/mosquito dynamics.

We evaluated existing methodology as well as our method using pooled Ct values to estimate WNV infection prevalence from pooled testing. Both methods yielded similar prevalence estimates for the Nebraska and Colorado data. However, pooled Ct values can be used to accurately estimate WNV infection prevalence at all prevalence levels, regardless of the pool size or the number of pools. Although the current gold-standard method using binarised data performs well at prevalence values less than 0.15 using 100 pools of 50 mosquitoes, it is unable to generate estimates at higher prevalences due to the saturation of positive pools. Therefore, the Ct-based approach may be more suitable under specific epidemiological conditions where underlying prevalence may exceed 10%, which can occur during focal outbreaks and more commonly towards the end of transmission season, where infection rates are high while mosquito abundance is declining. [52, 53, 54]. Additionally, the Ct-based method can provide more accurate estimates of prevalence when trapping effort is sparse. Using the binarised MLE approach, at least one pool in the catchment area must be negative WNV to calculate a prevalence estimate, which may not happen when few traps are used to conduct entomoloigcal surveillance. Furthermore, increasing both pool size and the number of pools improves accuracy with the Ct-based method (see Figure S20), but pool size does influence estimate accuracy with the binary-based method. As shown in Figure 6 and Figure S21, the maximum identifiable prevalence with binarised data decreases as the pool size increases. Another advantage of our Ct-value based method was that it allowed us to estimate the proportion and population-level prevalence of productive infections in a population, as opposed to all WNV-positive mosquitoes. This enables us to identify the proportion of infected mosquitoes which are infectious, which could be incorporated into more refined estimates of human risk.

Throughout this study, we based our inferences on synthetic Ct value distributions generated by our agent-based model, which leads to a number of limitations. While our estimates aligned well with those derived from binary data, we parametrised the viral kinetic models for both birds and mosquitoes using values from the literature [55], and more robust results would require data from deliberate infection and pooling experiments. For instance, experimentally deriving the distribution of Ct values for *k* positives in a pool of *n* would provide more realistic Ct value distributions than simulation alone. Our model also simplified the system by treating all mosquito and bird species interchangeably, assuming a single vector and a single host species. In reality, variation in viral kinetics between different species likely affect viral load kinetics, especially in regions where multiple vectors play a role in driving transmission. We attempted to parametrise the ecological aspects of the agent-based model using the literature. Still, we note that some parameters, particularly relating to overwintering, are not well defined. Finally, in this study, we did not conduct any formal model fitting to the data using our agent-based model. Our main objective of this study was not to formally fit the multi-scale agent-based model to the data, but rather to understand how individual-level viral kinetics in mosquitoes are reflected in pooled surveillance data, and to demonstrate the use of pooled Ct values in estimating WNV infection prevalence. However, it would be helpful to conduct formal model fitting, for example using an emulator [56], and then replicate the experiment to test the viral inheritance hypothesis to further solidify our results.

In this study, we did not model spatial structure nor estimate WNV infection prevalence at the county level, notably for Nebraska where surveillance data were collected across multiple counties and sites. All estimates should therefore be interpreted as averages across all trapping sites within each state. Although accounting for spatial heterogeneity may provide more locally relevant prevalence estimates, it is beyond the scope of this study. Additionally, we did not formally associate the prevalence estimates from either approach with human cases that occurred in Colorado and Nebraska during the study period. Our primary aim was to model possible biological mechanisms which could explain observed variation in pooled WNV Ct values accounting for within-host kinetics, and we therefore did not model spatial variability or associate estimated mosquito prevalence with human incidence.

There is a robust literature regarding the vector competence—the ability of a mosquito to transmit a pathogen—of WNV in relevant vector species; however, no studies have explicitly made the connection between Ct values, replication kinetics and infectiousness in mosquitoes. Exposing cohorts of mosquitoes to a range of viremias or allowing mosquitoes to feed on birds through a course of infection and taking daily measurements of virus replication, dissemination, and infection potential would provide high-resolution insights into viral kinetics in mosquitoes and the relationship between infectiousness and Ct values. Additionally, temperature affects EIP and needs to be considered when quantifying WNV growth kinetics through time in a laboratory setting [7, 28]. From our model, a key future direction for WNV surveillance is to better interpret the distribution of Ct values as a mixture of both infected and infectious mosquitoes. Population-level modelling, as we have done here, is a key step towards providing plausible mechanisms and frameworks for interpreting this variability, but deliberate laboratory-based infection experiments in mosquitoes to better understand the relationship between Ct values and infectivity will be necessary to confirm its impact on transmission dynamics.

WNV transmission, like other arboviruses, is heterogeneous in time and space, making timely entomological surveillance an invaluable tool for understanding transmission risk and informing control[16]. Increasing WNV prevalence in mosquito populations is a reliable indicator of increased human risk in the near future and is often used to initiate WNV control mechanisms such as spraying insecticides or asking the public to modify behaviours [54, 57, 58, 59]. Robust estimates of mosquito prevalence across a range of surveillance approaches and epidemiological scenarios are therefore crucial to inform public health policy. As arbovirus surveillance is routinely done via RT-qPCR, Ct values produced from these assays represent an untapped dataset to better understand transmission dynamics and inform human risk that comes at no additional cost to public health entities. This study represents the first effort to understand and model Ct value dynamics observed in WNV entomological surveillance and use these data to produce more refined estimates of WNV prevalence in mosquito populations. The results of this study indicate that viral kinetics in natural mosquito populations are not uniform and binary interpretations of WNV surveillance likely miscalculate mosquito infection rates and, therefore, estimates of human risk.

Although our main focus here is on WNV due to the robust, routine entomological surveillance systems that exist in the US, these ideas may be generalisable for surveillance of other arboviruses such as chikungunya, dengue, and Zika virus [60, 61] that use semi-quantitative viral load data, particularly through pooled testing. Our approach might be generalisable for other data sources, particularly next-generation surveillance approaches such as metagenomics, which allows sequencing microbial communities from environmental samples [62]. As these methods allow for quantification of the viral load contained in a sample [63, 64, 65, 66], it may be possible to extend our method as a tool to identify transmission processes within wildlife populations for multiple pathogens simultaneously.

Arboviruses are a persistent and growing threat to global public health. Due to climate change, globalisation, and human activities, the distribution of arbovirus vectors continues to expand [5]. For example, WNV was recently detected in the United Kingdom [67], and warmer temperatures in Europe have seen local outbreaks of dengue, chikungunya, WNV, yellow fever, and malaria, and it is predicted that they will continue to spread throughout Europe [68, 69]. Furthermore, as we have discussed and illustrated, there is still a large gap in the literature in understanding WNV (and other vector-borne diseases) ecology and biology. Vector-borne diseases are part of the complex ecological processes, and they are highly sensitive to environmental changes, non-human reservoirs, and human and non-human activities [70]. Therefore, an interdisciplinary and multi-scale approach to understand the interaction of between-and within-host biology of vector-borne diseases, environmental, and ecological processes is necessary to improve surveillance and optimise public health interventions.

## 5 Materials and Methods

### 5.1 Mosquito Collections and WNV Ct value Data

For both Colorado and Nebraska, *CO*_2_-baited CDC light traps are set at multiple sites to collect host-seeking mosquitoes. Data generated from gravid traps were not considered as our model assumes mosquitoes are collected while host seeking. Traps are set weekly during peak WNV transmission season from epidemiological weeks 22-39 (Nebraska) and weeks 23-37 (Colorado). Traps are placed for a single night, and collected mosquitoes are stored on dry ice, sorted by species, and placed into pools of up to 50 individual mosquitoes from the same trap prior to testing for WNV RNA via RT-qPCR [26]. Colorado mosquito surveillance data was provided by Colorado State University and generated as previously described [54]. Nebraska mosquito surveillance data was provided by the Nebraska Public Health Laboratory (NPHL). For identification purposes, *Cx. pipiens* species mosquitoes are not separated from the *Cx. salinarius* or *Cx. restuans*, which are nearly identical morphological. However, *Cx. pipiens* is far more abundant than either *Cx. salinarius* or *Cx. restuans* in Nebraska [71]. Mosquito pools were tested for WNV, St. Louis Encephalitis virus (SLE), and Western Equine Encephalitis virus (WEEV) via a trioplex RT-qPCR assay as previously described [27]. Notably, the same primer/probe sequences are used to detect WNV RNA in both the Colorado and Nebraska datasets. Following RT-qPCR, pools were reported as either positive or negative depending on the Ct value cut-off used (<40), alongside associated Ct values. Summary statistics from mosquito surveillance data for the Colorado and Nebraska datasets are reported in Supplementary Tables S2 and S3.

### 5.2 Modeling WNV Dynamics

We developed a mathematical model of WNV transmission between mosquitoes and birds and WNV viral dynamics, including within-host viral kinetics and simulated entomological surveillance to address our two study aims:

1. Explore potential mechanisms which might explain the time-varying signal in Ct values from pooled mosquito RT-qPCR testing.
2. Generate a synthetic dataset with which to develop and evaluate a novel estimator for WNV prevalence using Ct values from pools rather than binary positive/negative status.

We developed an agent-based model of mosquitoes and birds, inspired by the classic Ross-MacDonald model[31, 32], summarised in Figure 2. The model has three major components: 1) population-level interactions between *Culex*female mosquitoes and birds, 2) within-host viral dynamics of the mosquitoes and birds, and 3) routine weekly surveillance by capturing and pooling mosquitoes. All chosen parameter values are based on the literature, or, where necessary, based on informed assumptions (Supplementary Material S2).

#### 5.2.1 Population-level interactions between mosquitoes and birds, and other dynamics

In our model, we assumed that birds (hosts) may become susceptible (*S_H_*), infectious (*I_H_*), or recover (*R_H_*) with lifelong immunity following infection. We also model births and deaths of birds over the simulation, though these rates are slow relative to the rapid dynamics of the mosquito population.

We modelled aspects of the mosquito life cycle that directly influence the transmission cycle of WNV between birds and mosquitoes. In our model, mosquitoes are initially susceptible (*S_M_*) after birth and can become infected and infectious (*I_M_*) after biting an infected bird. Once a mosquito becomes infectious, it stays infected and infectious until its death (*D_M_*) or until it is captured. We assume that although there might be evidence for vertical transmission [72, 73], it is negligible. We consider only one mosquito species in the population with characteristics similar to *Culex pipiens* or *Culex tarsalis*. Transmission was assumed to occur with some probability following a bite between infected mosquitoes and susceptible birds and vice versa. Once a mosquito bites and takes a blood meal, we assume that it does not seek another blood meal for a number of days, assumed to be Poisson distributed with rate 4 day^−1^. In addition to infection states, we consider the two major stages of the mosquito life cycle: egg and adult. We assume that female mosquitoes lay eggs which hatch into adult female mosquitoes after an average of 8 days (see Figure 2). Male mosquitoes are not included in the model, as they do not contribute to the transmission cycle and are not represented in the observed data.

Mosquito biting, birth and death rates were assumed to be temperature dependent, which we proxied by varying these rates over the year with a sinusoidal function (see Supplementary Material S2) [74, 75, 76]. Furthermore, we assume that the adult mosquitoes will always either overwinter or actively contribute to transmission, depending on the environment’s temperature. Assumed parameter values are shown in Supplementary Material S2.

### 5.3 Within-host viral kinetics of birds and mosquitoes

The model incorporates an explicit viral kinetics—or Ct dynamics—framework for infected birds and mosquitoes. For birds, following infection, log viral load is assumed to increase linearly from the limit of detection immediately following infection until it reaches a peak a few days later (approximately 2.5 days). We then assume that the virus is cleared at a constant rate (on the log scale) over time. See Supplementary Material S3 for further details.

In contrast, we assume that mosquitoes can inherit the viral load of the bird they were infected by. In this model, the initial viral load of the infected mosquito is proportional to the viral load of the infected bird at the time of the bite. This reflects the possibility that the infectious dose in the infectee from a bite depends on the virus abundance in the infector. Once infected, mosquitoes can follow two possible fates: 1) the mosquito ingests detectable viral material from the bird but is not productively infected, and viral load remains static at the inherited value; 2) the mosquito becomes infected, and the viral kinetics model is initiated from the inherited viral load and reach the peak within days, and the number of days are normally distributed with mean 5 and standard deviation 0.1 (see Supplementary Material S4). For mosquitoes, infection is considered lifelong, with viral load remaining at its peak until the mosquito’s death.

Motivated by previous models of other pathogens (e.g., [77]), we assume that each individual’s model parameters are sampled from a normal distribution (see Supplementary Material S3 for the parameter values), ensuring individual-level variability in viral kinetics. See the Supplementary Material S2 for an illustration of the viral load dynamics of the birds and mosquitoes, the corresponding Ct dynamics models, and details about the parameters and their construction within a hierarchical structure in the Ct dynamics model.

### 5.4 Routine weekly surveillance through pooled RT-PCR testing of captured mosquitoes

We modelled weekly surveillance of WNV prevalence in trapped mosquitoes as a post-processing step, aiming to replicate routine entomological surveillance. After randomly sampling mosquitoes for each pool based on Nebraska data (see Supplementary Material S4 for further details), we calculate the pooled Ct value based on the methodology outlined by [78]. Here, the Ct values of each pool were transformed to the corresponding viral load and the average of the viral loads in the pool (assuming that negative mosquitoes contribution no viral load) were counted as the pooled viral load. Finally, this pooled viral load was transformed back to a Ct value (see Supplementary Material S4 for further details). We repeated these calculations for all the pools each week.

### 5.5 Estimation of WNV Prevalence in the Mosquito Population

We determined whether WNV prevalence in a mosquito population could be accurately estimated using the pooled Ct values obtained each week. Additionally, we examined whether prevalence estimates could be improved by incorporating Ct values, as opposed to relying solely on binarised positive/negative test results.

We calculated WNV prevalence using methodologies based on [19] and [79]. We constructed a Maximum Likelihood Estimator (MLE) by taking into consideration the conditional probability density distribution of an observed pooled Ct value given the possible number of positive samples in the pool. Details on how we calculated these density distributions and the corresponding MLE estimator are detailed in Supplementary Material S8. Point estimates for prevalence and confidence intervals were estimated using the MLE function from the *bbmle* package in R [80], or where the MLE function failed, a profile likelihood approach [81, 82].

We extended this method by jointly estimating the prevalence of non-productive and productive infections by maximising the likelihood of observed pooled Ct values across all pools. Each pool’s likelihood was obtained by summing over all possible combinations of productive, non-productive and negative samples in the pool, weighted by their multinomial probabilities and the empirical distributions of the Ct values (Supplementary Material S8).

To compare the accuracy of the estimates, we compared the estimated time-dependent prevalence with the true WNV prevalence in the mosquito population. We compared our Ct-based estimates to those obtained using binarised test results with the R package *PooledInfRate*, developed by the CDC [39]. This method generates a bias-corrected Maximum Likelihood Estimate (MLE) of WNV infection prevalence. The MLE is constructed based on the assumption that individuals in the pools follow independent and identically distributed Bernoulli distributions, and once tested, the pooled tests follow a Binomial distribution. See the Supplementary Material for further methodological details.

## 6 Acknowledgements

We would like to acknowledge Greg Ebel and Tobias Koch from Colorado State University and Ryan Vincent and Matt Parker from the City of Fort Collins for providing WNV surveillance data from Colorado used in this study. We would also like to acknowledge personnel at the Nebraska Public Health Laboratories and the Nebraska Department of Health and Human Services for providing WNV surveillance data from Nebraska used in this study.

## 8 Availability of codes and related data

All the code and data required to reproduce the analyses are available online at https://github.com/PunyaAlahakoon/west_nile_virus_abm.git.

