## supplementary_material for "Tracking West Nile virus dynamics using viral loads from trapped mosquitoes"

**S7 Model testing** **17**

**S9 Potential effects of pool size and number of pools on the accuracy of prevalence estimation** **24**

**S10 Availability of codes and related data** **31**

### S1 Nebraska and Colorado additional figures and data visualisations

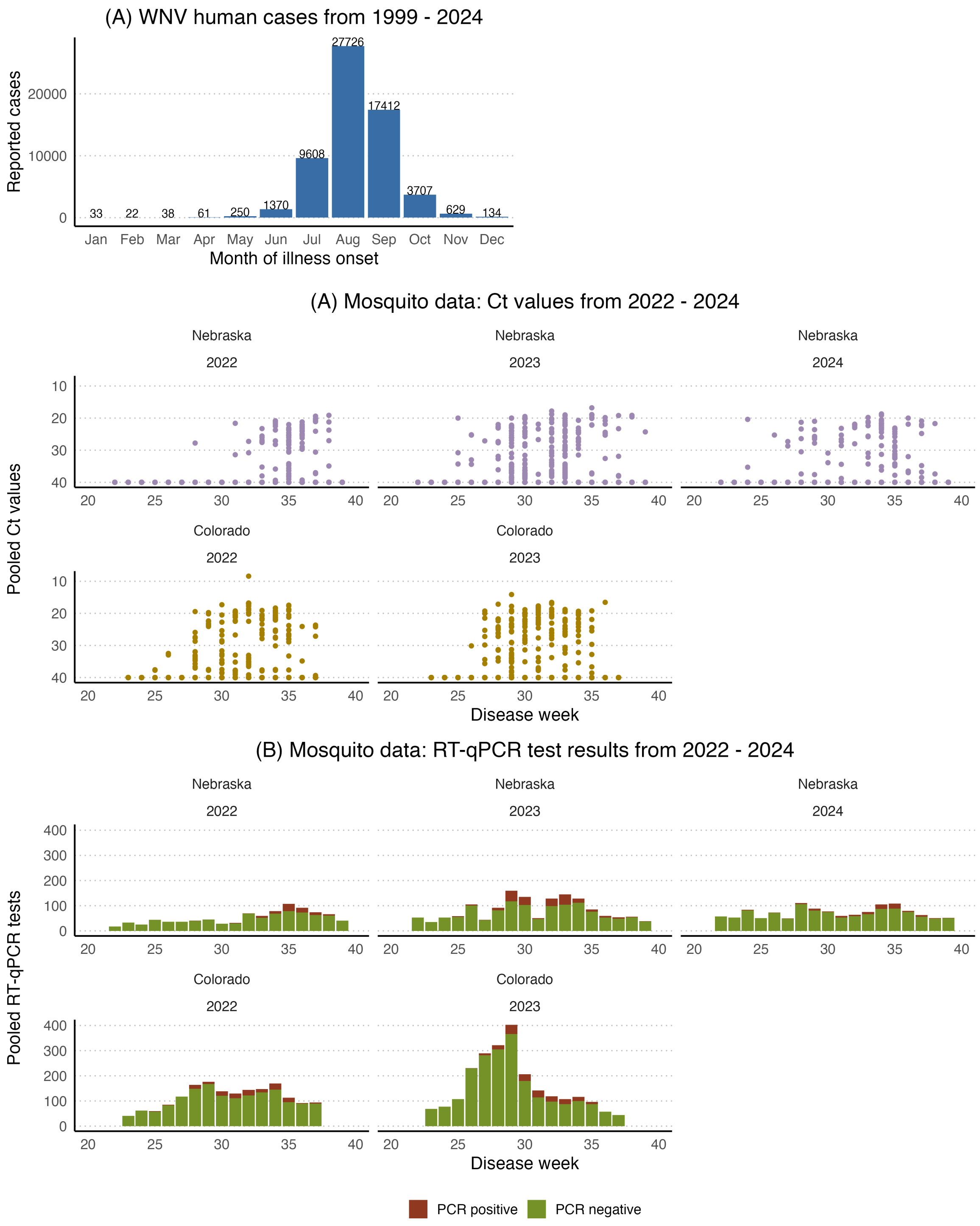

Figure S1: **Panel (A):** Observed monthly human cases from 1999-2024 (downloaded from the CDC website [1]).

**Panels (C) and (D)** [Figure 1 of the manuscript]: Observed data: Calculated pooled Ct value vs. disease week from 2022 to 2024. Pooled Ct values presented here are calculated by using species *Culex pipiens/restuans/salinarius* and *Culex tarsalis*.

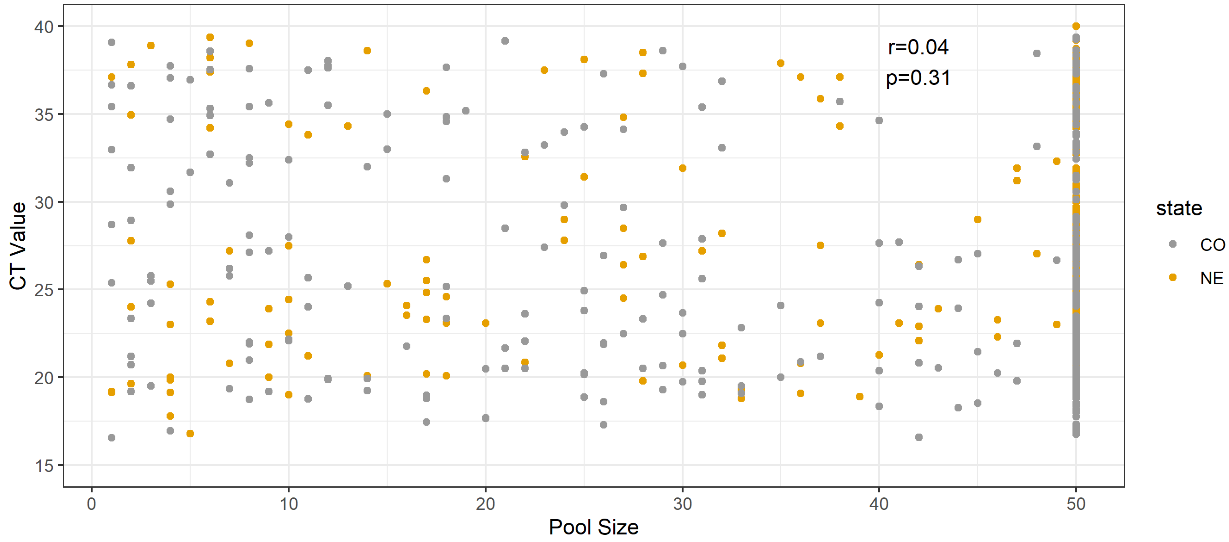

Figure S2: Observed Ct values distributed by number of mosquitoes contained in the tested pool. Pearson correlation coefficient = 0.04, p-value = 0.31.

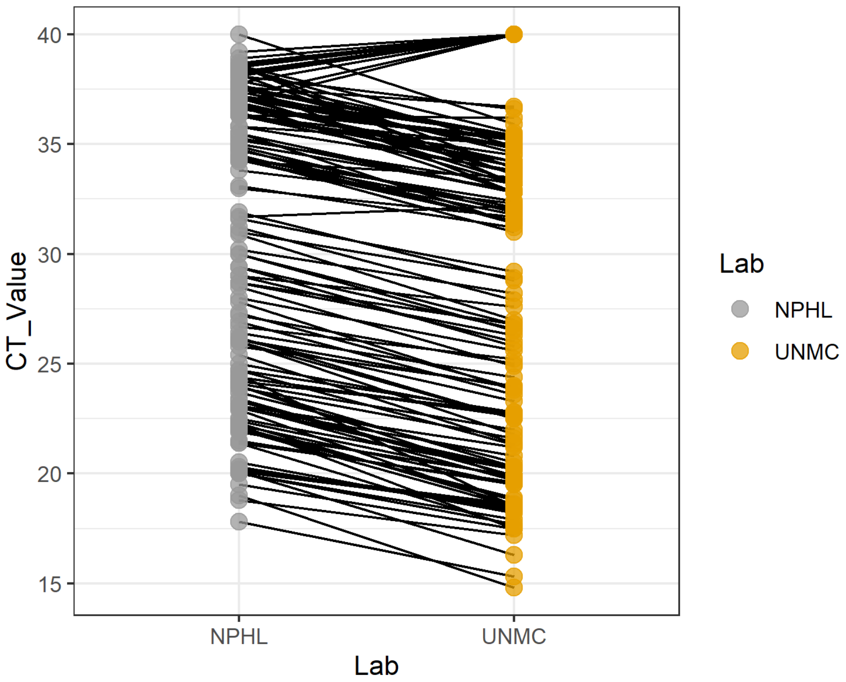

Figure S3: Observed Ct values from 143 samples tested in two laboratories. Ct values generated from the UNMC laboratory were consistently lower than those generated from NPHL, showing reproducibility of Ct value data from the same mosquito pool

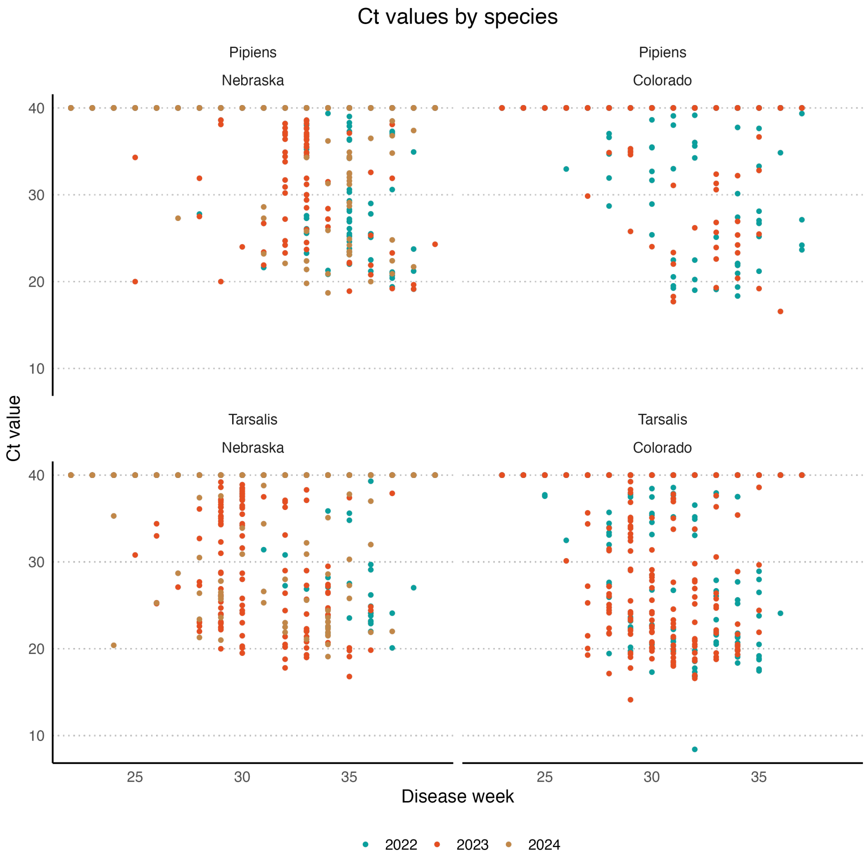

Figure S4: Ct values by species in Nebraska from 2022 to 2024 and Colorado from 2022 to 2023. Here, pipiens in Nebraska include *Culex pipiens/restuans/salinarius* species.

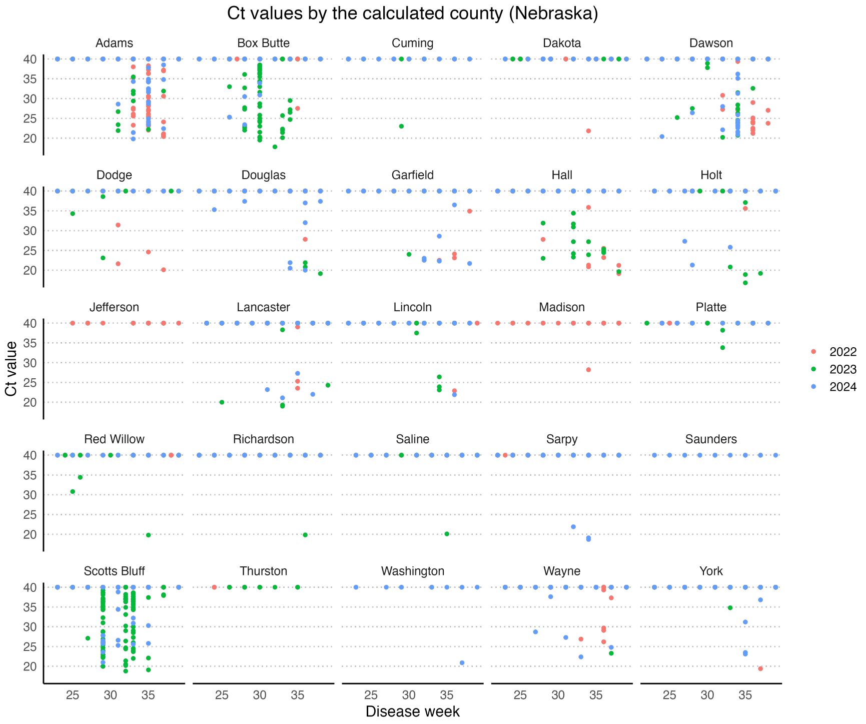

Figure S5: Ct values obtained by counties in Nebraska

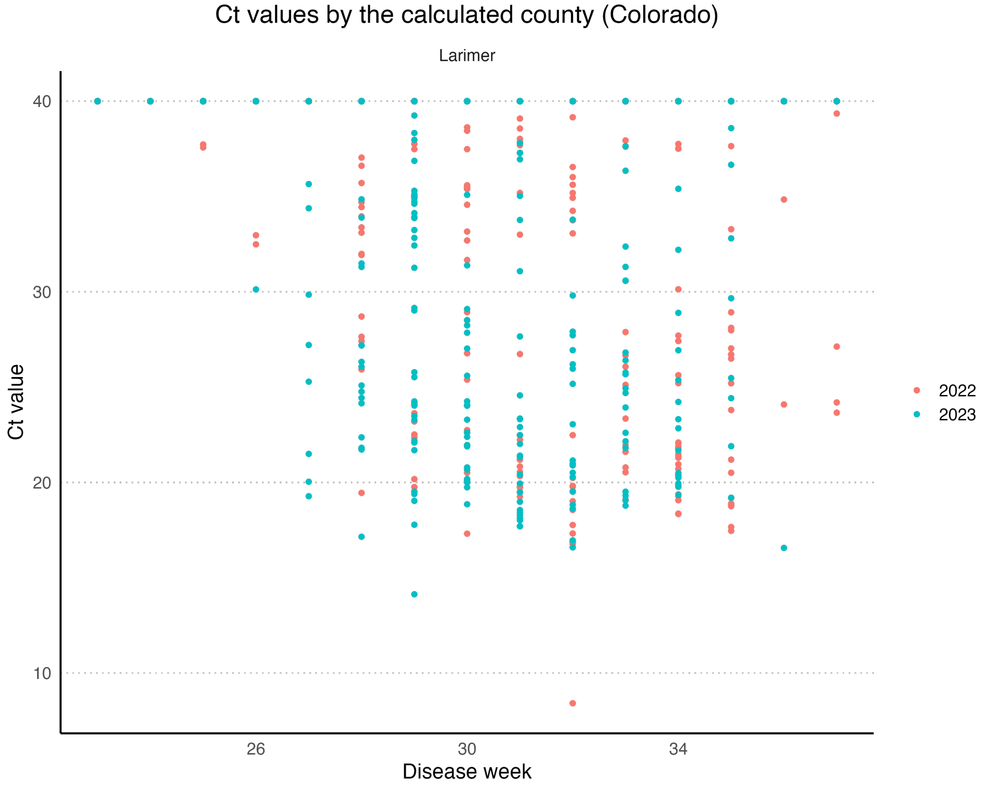

Figure S6: Ct values obtained by counties in Colorado.

#### S2 Population level parameters

Table S1: Population-level parameters of the agent-based model

| Parameter | Description | Values | Reference |
|---|---|---|---|
| $T$ | Simulation time | 26 years (excluding a burn-in period of one year) | NA |
| $t$ | Current time of the simulation (in days) | NA | NA |
| $N_H$ | Number of hosts(birds) at $t = 0$ (initial bird population size) | 1000 | NA |
| $N_M$ | Number of female mosquitoes at $t = 0$ (initial mosquito population size) | 10000 | female mosquito to bird ratio 1:10 [2] [3] |
| $I_{M0}$ | Infected female mosquitoes at $t = 0$ | 100 | NA |
| $p_{HI}$ | Transmission probability from mosquito to bird per bite | 0.88 | [4] |
| $b_{Mt}$ | Average number of eggs laid by mosquitoes who will eventually become adult female mosquitoes per day. | See details in Section S2.1 | |
| $a_{EA}$ | The number of days on average that takes for an egg to become an adult mosquito | 8 | CDC [5] |
| $b_H$ | Bird birth rate per day. | $0.023 \times N_H$ per day | 0.023: [6] |
| $d_H$ | Bird death rate | 0.0015 per day | [6] |
| $d_{Mt}$ | Daily death rate of mosquitoes | See details in Section S2.4 | |
| $p_{MIt}$ | Time-dependent transmission probability from bird to mosquitoes per bite | See details in Section S2.2 | |
| $r_t$ | Daily biting rate | See details in Section S2.3 | |
| $O_t$ | Overwintering period (time-dependent) | See details in Section S2.5 | |
| $p_{ot}$ | Probability overwintering (time-dependent) | See details in Section S2.5 | |
| $A_t$ | Mosquito active period (time-dependent) | See details in Section S2.6 | |
| $p_{at}$ | Probability of mosquitoes becoming active after overwintering (time-dependent) | See details in Section S2.6 | |

##### S2.1 $b_{Mt}$ calculation

According to [7], the per capita birth rate of mosquitoes is 0.537 per day, and the proportion of individuals who survive from egg to adult is 0.054. We calculated that the total number of eggs laid by mosquitoes that eventually become adults per day is $0.537 \times N_M \times 0.054 \approx 290$ on average.

We further assumed that $b_{Mt}$ depends on temperature (see [2] for more details). We modelled the temperature fluctuations using a cosine function (see Figure S7 for temperature-dependent birth rate for a year) with a mean rate of 290 and an amplitude of 1.

##### S2.2 Time-dependent transmission probability from bird to mosquitoes per bite, $p_{MIt}$

The probability of transmission from bird to mosquito per bite is assumed to be temperature-dependent. We assumed that the minimum probability, occurring during low-temperature periods (winter), is 0.04, the moderate temperature period is 0.09, and the maximum temperature period (summer) is 0.34 [2]. Accordingly, we calibrated a cosine function to reflect these factors. The function had the form average rate $\times (1 - \alpha \times cos(2\pi t/365))$, where $\alpha$ is the amplitude coefficient. See Figure S7 for the temperature-dependent probability from a bird to mosquitoes per bite. We use temperature and time-dependent interchangeability here, as changing one would result in a change to the other.

##### S2.3 Daily biting rate, $r_t$

Similar to $p_{MIt}$, we assume that the daily biting rate is also temperature dependent. Based on [2], we assume the biting rate during the low-temperature period to be 0.14, the moderate-temperature period rate to be 0.17 and the high temperature rate to be 0.2. Accordingly, we calibrated a cosine function to reflect these values. Figure S7 illustrates the temperature-dependent daily biting rate.

##### S2.4 Daily death rate of the mosquitoes, $d_{Mt}$

We assumed that the daily death rate of the mosquitoes is time-dependent and that the maximum number of mosquito deaths occurs when the temperature is at a maximum (that is the summer period). We used the maximum rate of death in the summertime to be 0.029 [6]. Accordingly, we calibrated a sinusoidal function to reflect this (see Figure S7).

##### S2.5 Overwintering period ($O_t$) and probability of overwintering $p_{ot}$

Building upon [8], we assume that the mosquito overwintering period starts in November and ends in February (see Figure S19). We further assume that the probability of overwintering changes over time (that is, sinusoidal) and the peak overwintering probability occurs in late December (see Figure S19).

##### S2.6 Mosquito active period and probability ($A_t$) of mosquitoes becoming active after overwintering ($p_{at}$)

Similar to the overwintering period, we assume that mosquitoes start becoming active in early March and decline by the end of October. We assumed that the change in the probability of mosquitoes becoming active over time is sinusoidal. See Figure S19 for more details.

##### S2.7 Blood seeking behaviour

We assume that immediately after a blood meal, mosquitoes go through a digestive period (the number of days for each mosquito is sampled from a Poisson distribution with rate 4). Once they go through this period, they join the mosquitoes that actively contribute to the transmission process.

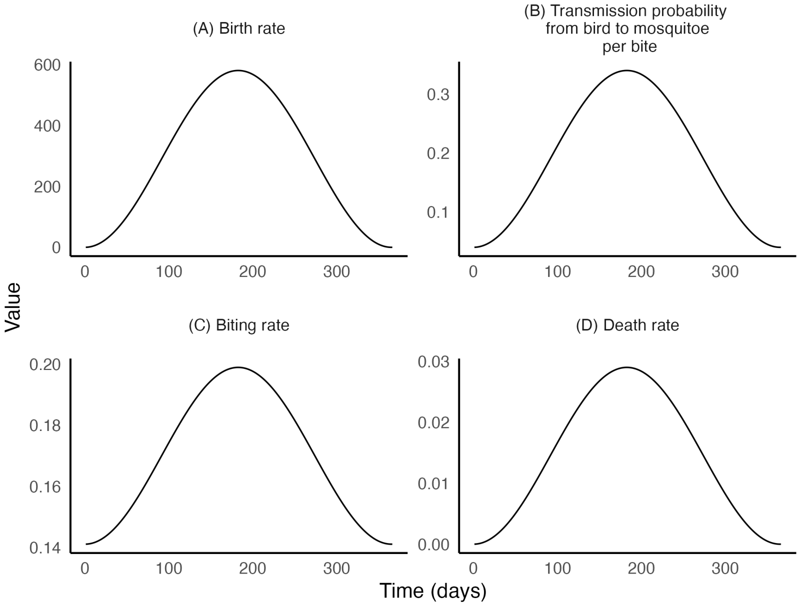

Figure S7: Time-dependent parameters for the mosquitoes for one year
**Panel (A):** Birth rate, $b_{Mt}$. **Panel (B):** Transmission probability from bird to mosquito, $p_{MIt}$. **Panel (C):** Biting rate, $r_t$. **Panel (D):** Daily death rate, $d_{Mt}$

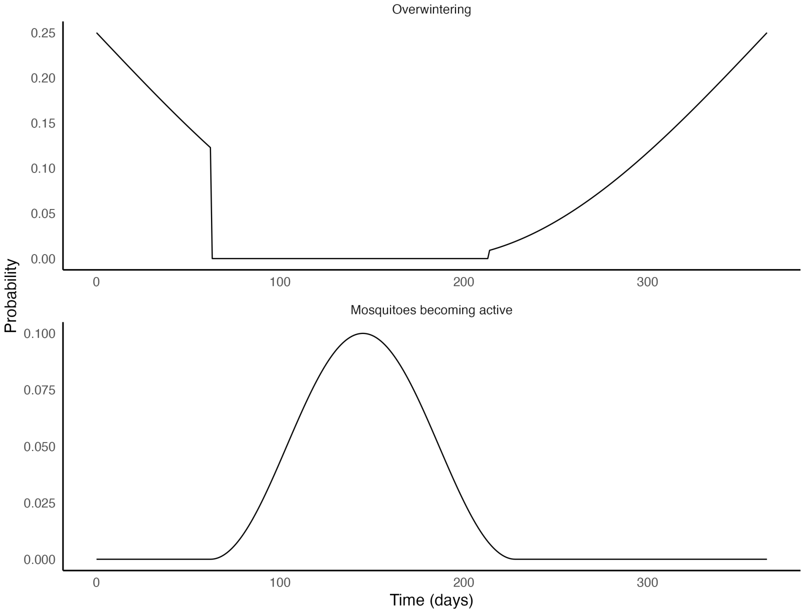

Figure S8: Probability of overwintering (top panel) and that of the mosquitoes becoming active (bottom panel) during one year period.

#### S3 Within-host dynamic parameters

The Ct dynamics models used in our agent-based model are presented in Figure S9. The top panel presents the Ct model for birds and the second panel Ct models used for the mosquitoes. For mosquitoes, the Ct dynamics models for the two assumptions ((a) and (b)) described in the manuscript are illustrated in the bottom panel. Generally, the increase or decrease of the Ct values is calculated by a $y = m \times t + y_0$ type relationship with $y$ indicating the Ct value at time $t$, $m$, the slope and $y_0$, the intercept.

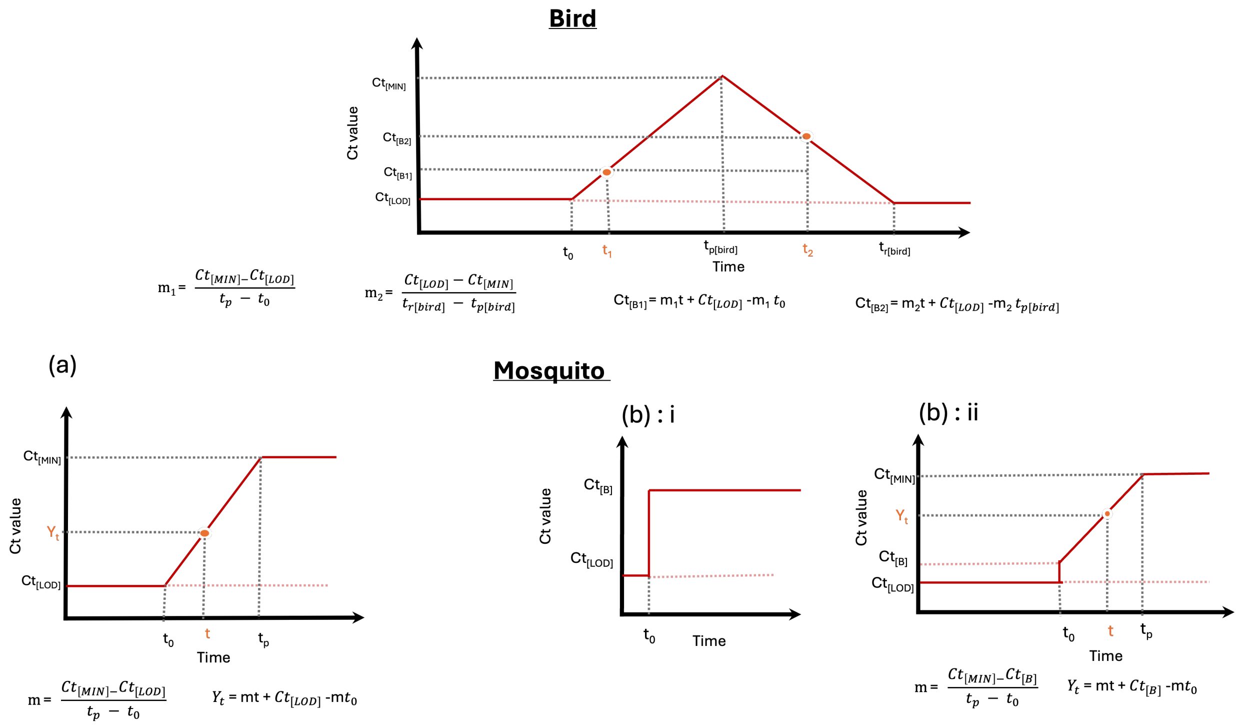

Figure S9: Ct dynamics models used for birds (top panel) and the mosquitoes (bottom panel) in our agent-based model.
**Top panel:**
$m_i$ (where $i = 1, 2$) represents the slope parameters for the birds. $C_{[Bi]}$ ($i = 1, 2$) represents a random Ct value within a bird since infection.
**Bottom panel**:
**Panel (a):** Ct dynamics model when the mosquitoes undergo dynamic infection from $Ct_{[LOD]}$, the limit of detection (which is 40 in our study). $m$ represents the slope parameter of a mosquito in the Ct model. $y_t$ represents a random Ct value of a mosquito since infection.
**Panel (b):** Mosquitoes inherit a certain percentage of viral load from birds. (i) shows the static viral load, and (ii) illustrates the dynamic viral load models. $Ct_{[B]}$ represents the Ct value inherited from the bird at the time of a successful infection of a mosquito.
For both bird and mosquito Ct dynamics models, $Ct_{[LOD]}$ represents the limit of detection (Ct=40), $t_0$ is the time of infection, and $Ct_{[MIN]}$ is the minimum Ct value that reaches the peak viral load.

In our agent-based model, we assume individual-level heterogeneity in Ct dynamics in birds and mosquitoes. That is, the parameters that enable the calculation of slopes and intercept terms in the Ct value dynamics model, are different. For birds, these parameters are time for peak viral load ($t_{p[bird]}$), minimum Ct value at peak viral load ($Ct_{[MIN]}$), time for viral load clearance and the Ct value comes to the limit of detection, $t_{r[bird]}$. For mosquitoes, these parameters are time for peak viral load ($t_p$), and the minimum Ct value at peak viral load ($Ct_{[MIN]}$).

For the bird and mosquito agents in the agent-based model, we sampled parameters randomly from the following distributions.

For birds:

$$t_{p[bird]} \sim Normal(2.5, 0.1^2) \tag{S.1}$$
$$Ct_{[LOD]} - Ct_{[MIN]} \sim Normal(15, 0.1^2) \tag{S.2}$$
$$t_{r[bird]} \sim Normal(5, 0.3^2) \tag{S.3}$$

Therefore, for birds, Ct value at time $t$, $Y_{t[bird]}$ is,

$$Y_{t[bird]} = \begin{cases} Ct_{[LOD]}, & \text{if } t \leq t_0 \\ -\frac{(Ct_{[LOD]}-Ct_{[MIN]})}{(t_0-t_{p[bird]})}t + Ct_{[LOD]} + \frac{(Ct_{[LOD]}-Ct_{[MIN]})}{(t_0-t_{p[bird]})}t_0, & \text{if } t_0 < t \leq t_p[bird] \\ \frac{(Ct_{[LOD]}-Ct_{[MIN]})}{(t_{r[bird]}-t_{p[bird]})}t + Ct_{[MIN]} - \frac{(Ct_{[LOD]}-Ct_{[MIN]})}{(t_{r[bird]}-t_{p[bird]})}t_{p[bird]}, & \text{if } t_p[bird] < t \leq t_{r[bird]} \\ Ct_{[LOD]}, & \text{if } t \geq t_{r[bird]}. \end{cases} \tag{S.4}$$

For mosquitoes:

$$t_p \sim Normal(5, 0.1^2) \tag{S.5}$$
$$Ct_{[LOD]} - Ct_{[MIN]} \sim Normal(15, 0.2^2) \tag{S.6}$$
$$\tag{S.7}$$

Therefore, for mosquitoes, Ct value at time $t$, $Y_t$, under model (a), is

$$Y_t = \begin{cases} Ct_{[LOD]}, & \text{if } t \leq t_0 \\ -\frac{(Ct_{[LOD]}-Ct_{[MIN]})}{(t_0-t_p)}t + Ct_{[LOD]} + \frac{(Ct_{[LOD]}-Ct_{[MIN]})}{(t_0-t_p)}t_0, & \text{if } t_0 < t < t_p \\ Ct_{[MIN]} & \text{if } t \geq t. \end{cases} \tag{S.8}$$

For mosquitoes, Ct value at time $t$, $Y_t$, under model (b) : i, is

$$Y_t = \begin{cases} Ct_{[LOD]}, & \text{if } t \leq t_0 \\ Ct_{[B]} & \text{if } t > t_0, \end{cases} \tag{S.9}$$

where $Ct_{[B]}$ is the Ct value of the bird at the time of bite and infection.

For mosquitoes, Ct value at time $t$, $Y_t$, under model (b) : ii, is

$$Y_t = \begin{cases} Ct_{[LOD]}, & \text{if } t \leq t_0 \\ -\frac{(Ct_{[B]}-Ct_{[MIN]})}{(t_0-t_p)}t + Ct_{[B]} + \frac{(Ct_{[B]}-Ct_{[MIN]})}{(t_0-t_p)}t_0, & \text{if } t_0 < t < t_p \\ Ct_{[MIN]} & \text{if } t \geq t_p. \end{cases} \tag{S.10}$$

#### S4 Simulating mosquito trapping and pooling for routine surveillance

Mosquito trapping and the pooling stages were simulated as post-processing steps in our agent-based model.

For each week, we simulated the number and size of each pool by considering the data from Nebraska, United States. For week $x$, we first took the number of pools collected in Nebraska ($b_x$) and generated a random number ($r_{b_x}$) from a Poisson distribution with ($b_x$) as the rate parameter. Then we sampled $r_{b_x}$ number of pool sizes with replacement ($\boldsymbol{n_x}$) from the sample sizes observed in Nebraska on week $x$. Therefore, $r_{b_x}$ and ($\boldsymbol{n_x}$) were considered the number of pools and the pool samples generated in week $x$, respectively, that replicated the mosquito capture behaviour. Let the total number of mosquitoes in all pools be $N$.

Next, on week $x$, we considered the available mosquitoes (susceptible or infected) that were simulated. Of them, we sampled mosquitoes $N$ without replacement and assigned them to the pools. However, if the number of mosquitoes available from the simulation is less than those that were meant to be captured, we re-adjusted $N$ by reducing the number of pools in week $x$ so that $N$ is less than or equal to the available simulated mosquitoes in week $x$. We repeated this procedure for all the disease weeks.

##### S4.1 Calculating Ct values from pooled test

First, we calculated the underlying viral load of the captured mosquitoes by considering the following log-linear relationship between the viral loads and $Ct$ values (see Figure S10 for the relationship).

$$v_i = 10^{(y-y_0)/m}, \tag{S.11}$$

where $v_i$ is the viral load, $y_0$ is the intercept term, $m$ is the slope, and $y$ is the observed Ct value. Here, $y_0$ and $m$ are calculated from the data by fitting Ct values to a standard curve of known WNV RNA quantity in a 10-fold serial dilution through a regression analysis. We assume that mosquitoes that are not infected will have a viral load of 0 copies/ml, corresponding to a Ct value of 40. Here, $y_0 = 36.9$ and $m = -2.7$.

We calculate the overall viral load in the pool as the mean of the viral loads in the sample ($\bar{\nu}$). Then we calculated the Ct value of the pool with transformation (S.11). We repeat this procedure across all the weeks.

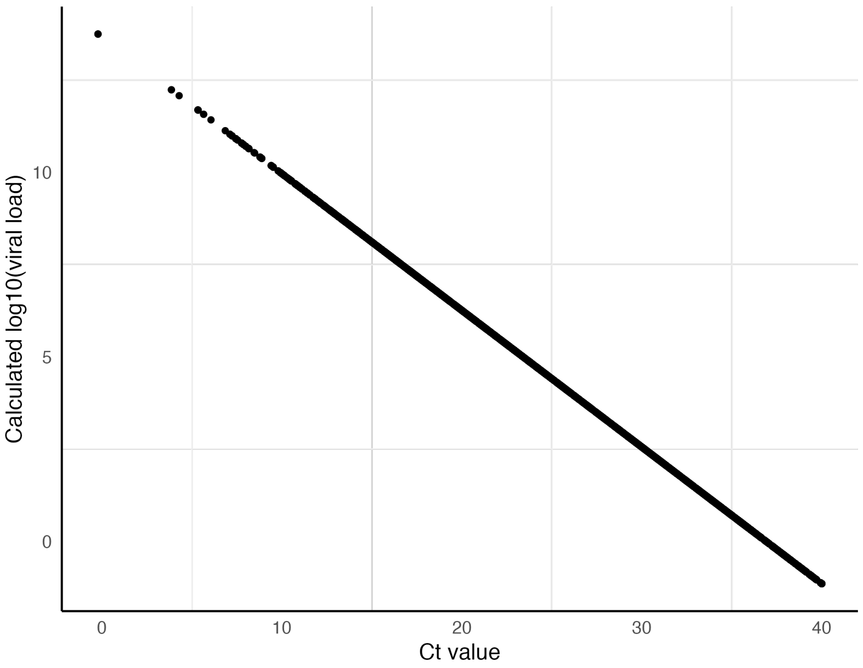

Figure S10: Relationship between log10(viral load) and Ct values.

#### S5 Pseudocode of the agent-based model

Given that the biting rate on day $t$ is $r_t$, we calculate the probability of a daily bite to be $(1 - e^{-r_t})$. Let a mosquito agent be denoted as $M_i$ for $i = 1, 2, \ldots, N_{Mt}$ and a bird agent be denoted as $B_j$ for $j = 1, 2, \ldots, N_{tH}$. Here, $N_{Mt}$ and $N_{Ht}$ are mosquito and bird population sizes at time $t$, respectively. Furthermore, let $M_{Ot}$ be the overwintering population at $t$ and $E_t$ be the egg population at time $t$.

The pseudocode of the agent-based model is summarised in the algorithm below.

**Initialization** Define and parametrise all the parameters defined in Table S1

**While** $t < T + 1$**,**

1. **Event:** Mosquito bite
   - (a) For $i = 1, 2, \ldots, N_{Mt}$ mosquitoes,
     - i. Generate a random number $r_1 \sim$ uniform(0,1)
     - ii. If $r_1 <= (1 - e^{-r_t})$, $M_i$ bites a bird. Randomly sample a bird $B_j$ from the population of bird agents.
     - iii. If $M_i$ is infected and $B_j$ is susceptible, generate $r_2 \sim$ uniform(0,1)
       - A. If $r_2 <= p_{HI}$, $B_j$ becomes infected.
     - iv. If $M_i$ is susceptible and $B_j$ is infected, generate $r_3 \sim$ uniform(0,1)
       - A. If $r_3 <= p_{MIt}$, $M_i$ becomes infected.
       - B. Store $B_j$'s viral load.
       - C. Store $M_i$ as "digesting" and set the number of days (generate $r_{di} \sim Poisson(4) + t$) until the next day of possibly taking a blood meal.
2. **Event:** Mosquito births
   - (a) Generate a random number of eggs, $R_e \sim$ Poisson($b_{Mt}$) and send $r_e$ eggs to an egg state.
   - (b) For all $k = 1, 2, \ldots, R_e$,
     - i. Generate $r_k \sim$ Poisson($a_{EA}$) $+ t_{i_{birth}}$, where $t_{i_{birth}}$ is the time the egg was laid.
     - ii. If $r_k >= t$, change the $k$th egg to an adult. Remove the egg from the egg population and add it to the adult mosquito population.
3. **Event:** Overwintering.
   - (a) If $t \subset O_t$, for $i = 1, 2, \ldots, N_{Mt}$ mosquitoes,
     - i. Generate $r_t \sim$ uniform(0,1).
     - ii. If $r_t < p_{ot}$,
     - iii. $M_i$ overwinters and does not contribute to the transmission cycle.
4. **Event:** Mosquitoes becoming active.
   - (a) If $t \subset A_t$, for $i = 1, 2, \ldots, N_{Mt}$ mosquitoes,
     - i. Generate $r_t \sim$ uniform(0,1).
     - ii. If $r_t < p_{at}$,
     - iii. $M_i$ becomes active and contributes to the transmission cycle.
   - (b) $M_i$ is in the "digesting" period,
     - i. If $r_{di} >= t$,
     - ii. $M_i$ starts seeking blood meals and contributes to the transmission cycle.
5. **Event:** Mosquito deaths
   - (a) For $i = 1, 2, \ldots, N_{Mt}$ mosquitoes
     - i. Generate $r_i \sim$ uniform(0,1)
     - ii. If $r_i <= (1 - e^{-d_{Mt}})$, $M_i$ dies.
     - iii. Remove $M_i$ from the mosquito agent population.
6. **Event:** Host recovery.
   - (a) For $j = 1, 2, \ldots, N_{Ht}$ birds,
     - i. If $B_j$ is infected, the current Ct value is at the limit of detection, and the time since infection is at least 4 days, $B_j$ recovers.

7. **Event:** Host deaths.
   (a) For $j = 1, 2, \ldots, N_{Ht}$ birds,
      i. Generate $r_j \sim$ uniform(0,1)
      ii. If $r_j <= d_H$, $B_j$ dies.
      iii. Remove $B_j$ from the bird agent population.
8. **Event:** Birth of hosts.
   (a) For $j = 1, 2, \ldots, N_{Ht}$ birds,
      i. Generate $r_j \sim$ Poisson($b_H$)
      ii. Add $r_j$ susceptible birds to the bird agent population.
9. **Store values at $t$:**
   (a) Infection state of all the mosquitoes.
   (b) Number of new mosquito eggs, and number of eggs that become adults.
   (c) Number of mosquito deaths.
   (d) Number of bites of each mosquito.
   (e) Infection state of all the hosts.
   (f) Number of host births and host deaths.
   (g) Ct values of the mosquitoes and the hosts.
   (h) Ct value parameters of mosquitoes and hosts.

### S6 Simulations from the agent based model

#### S6.1 Simulations of birds over 10 years, including the first year

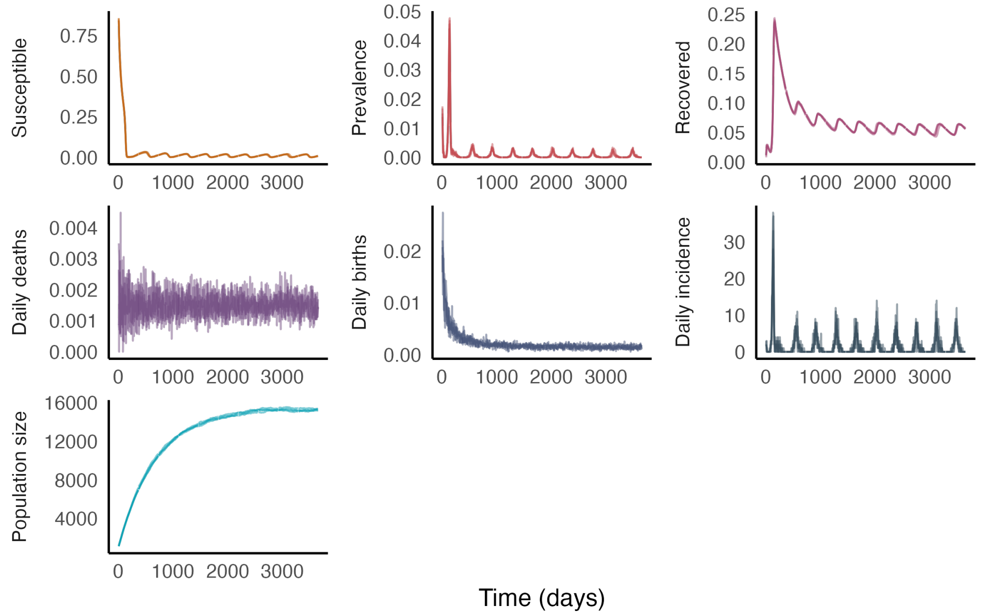

Figure S11: States of the hosts over ten years.

#### S6.2 Simulations of mosquitoes over 10 years, including the first year

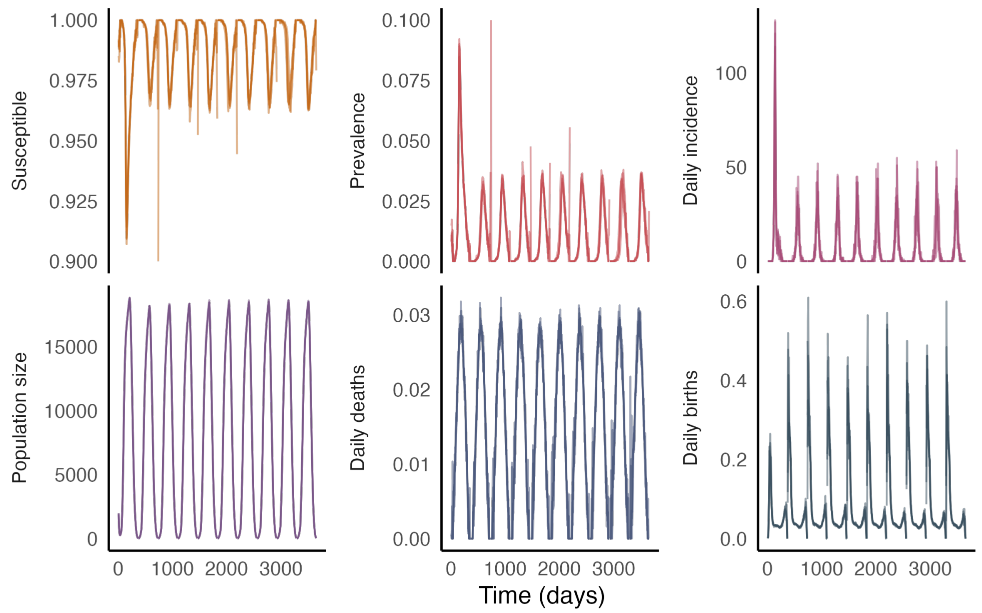

Figure S12: States of mosquitoes over ten years.

##### S6.3 Ct dynamics of birds

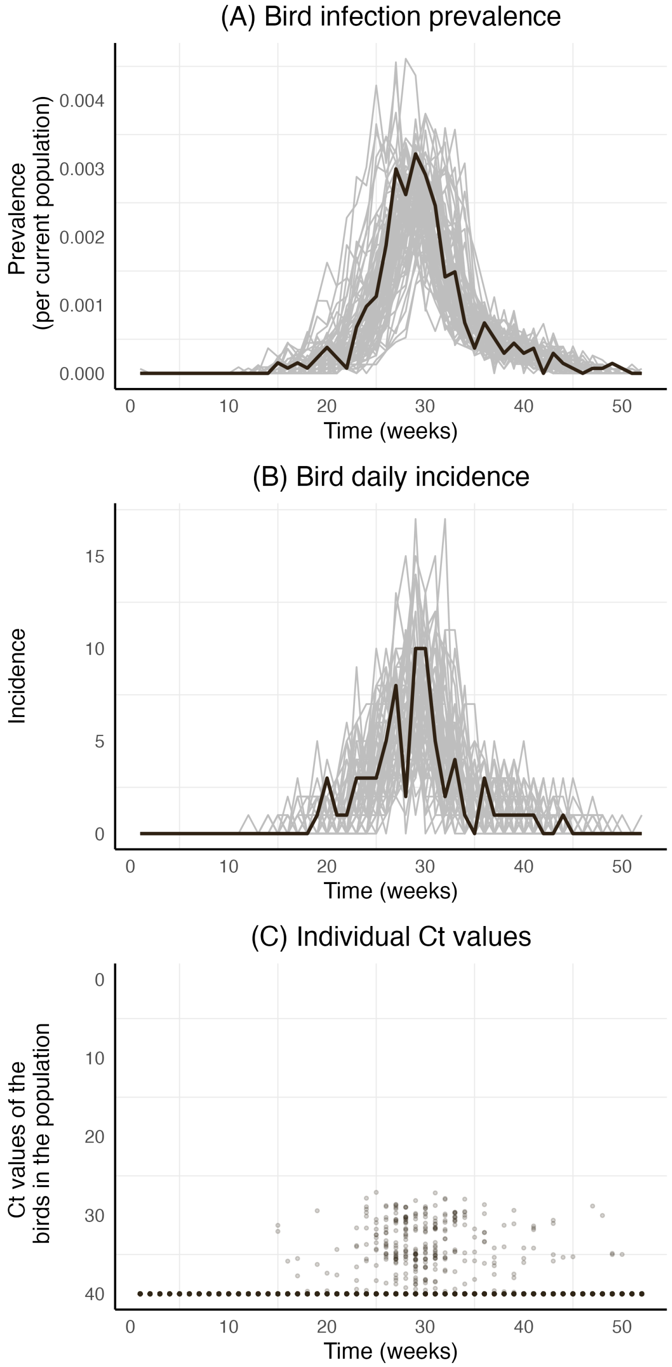

Figure S13: Simulated population-level WNV infection dynamics in mosquitoes. Each line represents one year of simulated data, and the bold black line highlights the simulated trajectory used for Panel (C) and Figure 4 in the manuscript.
**Panel (A):** Weekly bird infection prevalence per capita.
**Panel (B):** Weekly bird infection incidence per capita.
**Panel (C):** Individual-level Ct values of the bird population from one year of a single simulation. Each point represents a Ct value from a single randomly selected bird, representing perfect observations of the entire population at all times.

#### S7 Model testing

##### S7.1 Testing the model assumptions on the independence of the mosquito viral load dynamics from the bird viral loads at the time of infection and bite

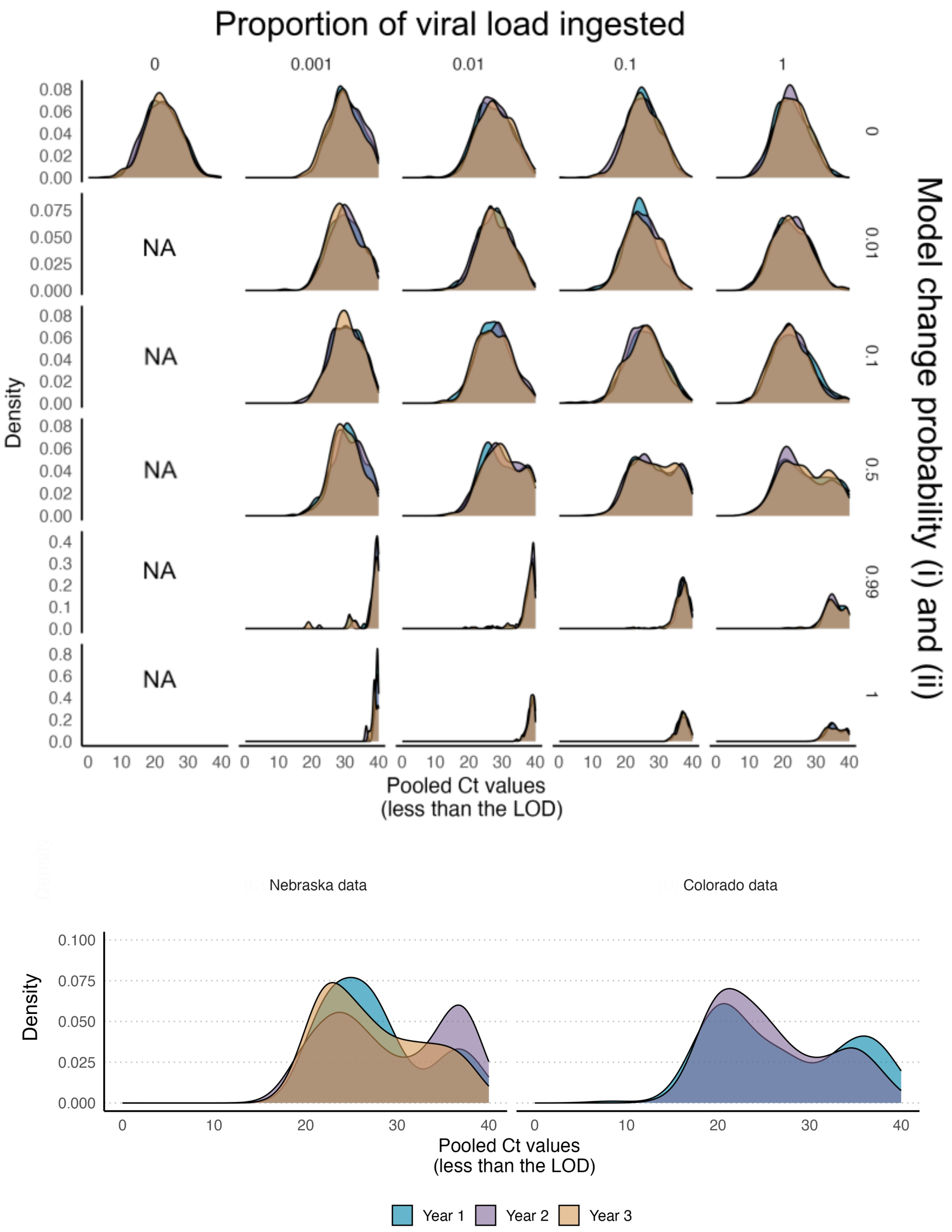

Figure S14: **Top panel:** Density distributions of the Ct values within the level of detection with change of viral inheritance probability vs. model change probability. Each density simulated per year is overlaid. Model change for mosquitoes was done as a Bernoulli trial with the probability specified in each sup-panel in the right corner. A successful Bernoulli trial indicated model (b) i: that is, the mosquito viral load will remain constant after successful viral inheritance.
**Bottom panel:** Density distributions of the Ct values within the level of detection for the Colorado and Nebraska datasets. All the densities for the three years are overlaid.

##### S7.2 Is a decline in viral load over time biologically plausible in mosquitoes?

In order to investigate whether decline in viral load over time is biologically plausible in mosquitoes, we ran the agent-based model with varying decay rates for the mosquitoes. We kept the proportions of productive and non-productive infection at 0.5 and the percentage of viral ingestion from the birds to be 100% as we observed this setting yielded Ct value distributions similar to those observed in Nebraska and Colorado (see Figure S14). We considered four different viral load decay rates for the mosquitoes after they become infected. These rates are 0.001, 0.01, 0.1, and 0.5. The Ct value distributions are illustrated in Figure S15 (left panel) with their corresponding Ct value dynamics (right panel).

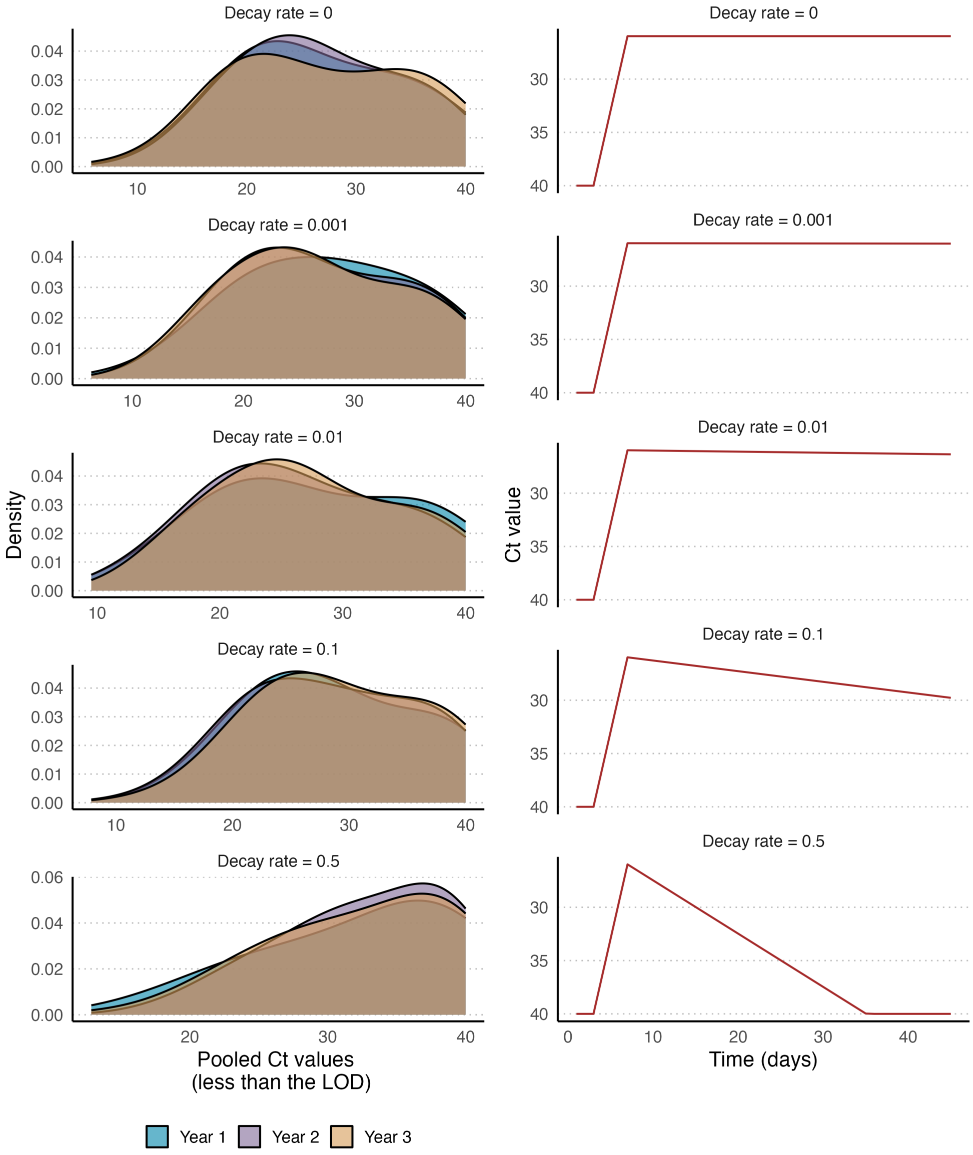

Figure S15: **Left** Density distributions of the Ct values within the level of detection with change of viral inheritance probability =1 vs. model change probability=0.5 with assumed decay rates 0, 0.001,0.01, 0.1, and 0.5. Each density simulated per year is overlaid.
**Right:** Ct dynamics with assumed decay rates 0, 0.001,0.01, 0.1, and 0.5.

#### S7.3 Models of mosquito within-host dynamics after 100% viral inheritance from birds

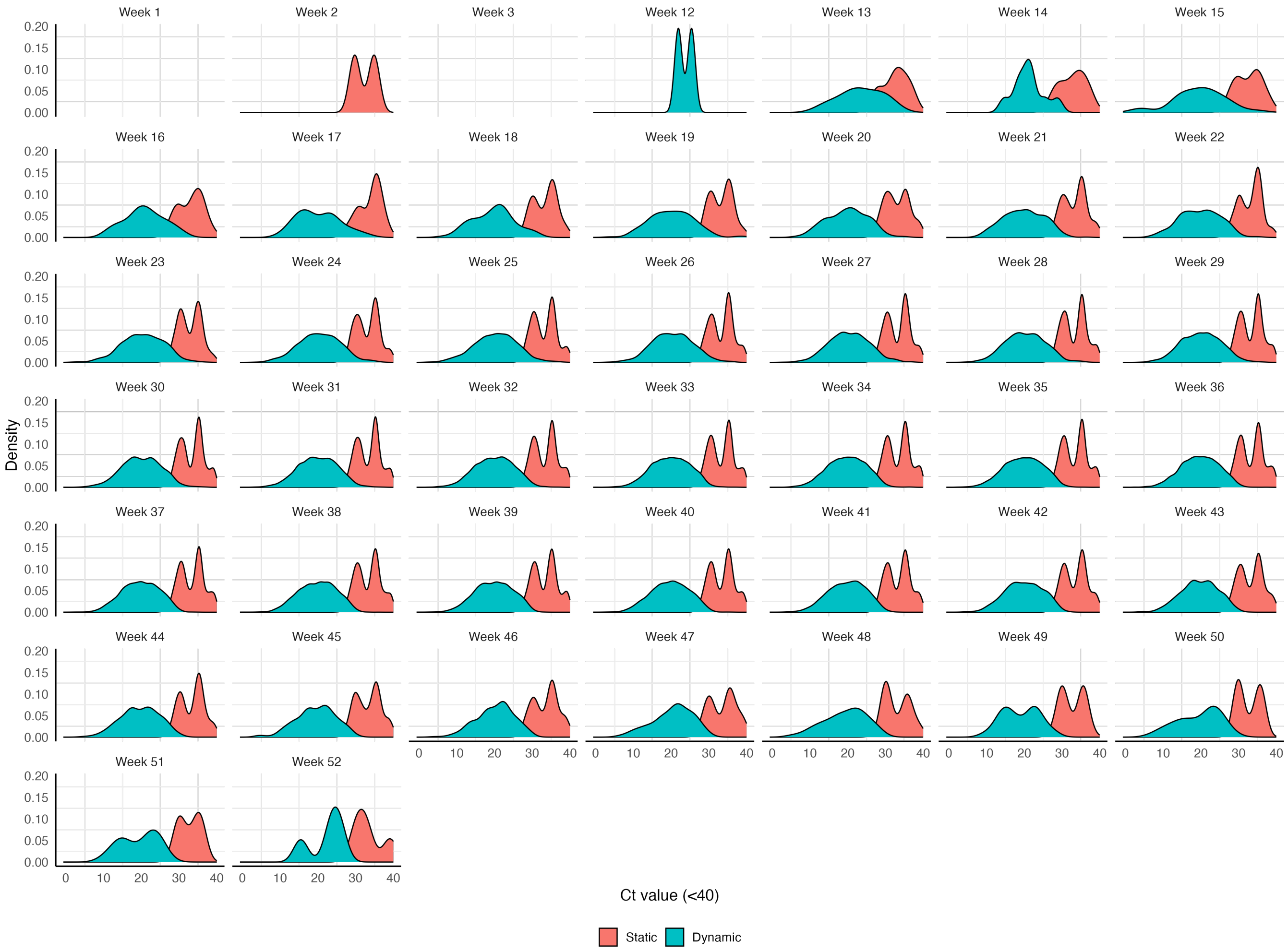

Figure S16: Density distributions of Ct values of the mosquitoes (less than the limit of detection) from simulated data assuming mosquitoes inherit 100% of viral load from birds, and 50% of mosquitoes undergo a static viral kinetics trajectory. Each panel illustrates the distributions of the Ct values of the mosquitoes that underwent a static (in pink) or dynamic (in blue) viral kinetics trajectory in each week.

#### S8 Maximum Likelihood Estimation (MLE) method to estimate the prevalence

In this study, we used the MLE method to estimate the prevalence of the WNV in the mosquito population. We first constructed a maximum likelihood function based on the methods by [9] and [10].

We first computed conditional probability densities for an observed pooled Ct value in a pool given that there are $k$ positive samples and considering all the possible values of $k$ (Figure S11&12). This method is similar to the method used in [9]. Let this conditional density be denoted as $Pr(y_i|k,n)$ where $y_i, k$, and $n$ represent the observed pooled Ct value, the number of positive samples and the pool size, respectively. The pseudo-code for calculating this conditional probability density is below.

For k from 0 to n;

1. Sample 10 0000 viral loads $v_{ki} = \sum_{j=1}^{k} z_j$ where $z_j$ are viral loads sampled from data (simulated from the agent-based model and filter the mosquitoes that were infected) without replacement. Here $i = 1, 2, \ldots, 10000$.
2. Calculate the mean viral load, $\bar{v_{ki}} = v_{ki}/n$ for all $i = 1, 2, \ldots, 10000$.
3. Calculate the pooled Ct value, $y_{ki}$ from the transformation, $y_{ki} = -2.7 \times log10(\bar{v_{ki}}) + 36.9$ for all $i = 1, 2, \ldots, 10000$.
4. If $y_{ki} > 40$, $y_i = 40$.
5. Estimate the kernel density (KDE) $\mathscr{K}_k$ from the $y_{ki}$ using the *adaptiveKernel* function in the *car* package in R with a bandwidth parameter 0.1.
6. Interpolate $\mathscr{K}_k$ within the 0-40 range of pooled Ct values.
7. Account for the false positive rate, $r$, if the number of samples in a pool is 1 to account for the Ct values that might arise from false positives. $r = 0.00001$, if the pooled Ct value is less than the limit of detection (40), otherwise, $1 - r$. $r$ is set to be arbitrarily small, as we do not expect false positives, but it is necessary to ensure a tractable likelihood surface.

Accordingly, the MLE estimate, $\hat{p}$ following from [10], was calculates as,

$$\hat{p} = \arg\max_{p} \prod_{j=1}^{b} \sum_{k_j=0}^{n_j} \binom{n_j}{k_j} p^{k_j}(1-p)^{(n_j-k_j)} Pr(y_j|k_j, n_j), \tag{S.12}$$

where $Pr(y_j|k_j, n_j)$ was approximated from $\mathscr{K}_k$. The approximated KDEs are illustrated in Figures S17 and S18. We used the *mle2* function in *bbmle* R package to calculate the prevalence estimate and the corresponding 95% confidence intervals. On occasions when the *mle2* function fails to estimate the prevalence, we used the profile likelihood method to estimate the confidence intervals.

When the productive and non-productive infections are taken into consideration, the MLE estimates for productive ($p_d$) and non-productive ($p_s$) infection prevalence were calculated as,

$$(\hat{p_s}, \hat{p_d}) = \arg\max_{(p_s,p_d)} \prod_{j=1}^{b} \sum_{k_j=0}^{n_j} \sum_{k_{jd}=0}^{k_j} \frac{n_j!}{k_{jd}!(k_j-k_{jd})!(n_j-k_j)!} p_d^{k_{jd}} {p_s}^{(k_j-k_{jd})} p_n^{(n_j-k_j)} Pr(y_j|k_j, n_j, k_{jd}), \tag{S.13}$$

where $Pr(y_j|k_j, n_j, k_{jd})$ is the joint KDE distribution for the observed pooled Ct values given the productive and non-infections, and $p_n(= 1 - p_s - p_d)$ is the prevalence of the negative samples. $k_{jd}$ is the number of productively infected samples.

To calculate the confidence intervals based on the profile likelihood method, we used a sequence of values for the negative log likelihood of the equation (S.13) given the sequence of possible values for the prevalence. Assuming a chi-squared distribution with 1 degrees of freedom for these values, we calculated the 95% confidence intervals.

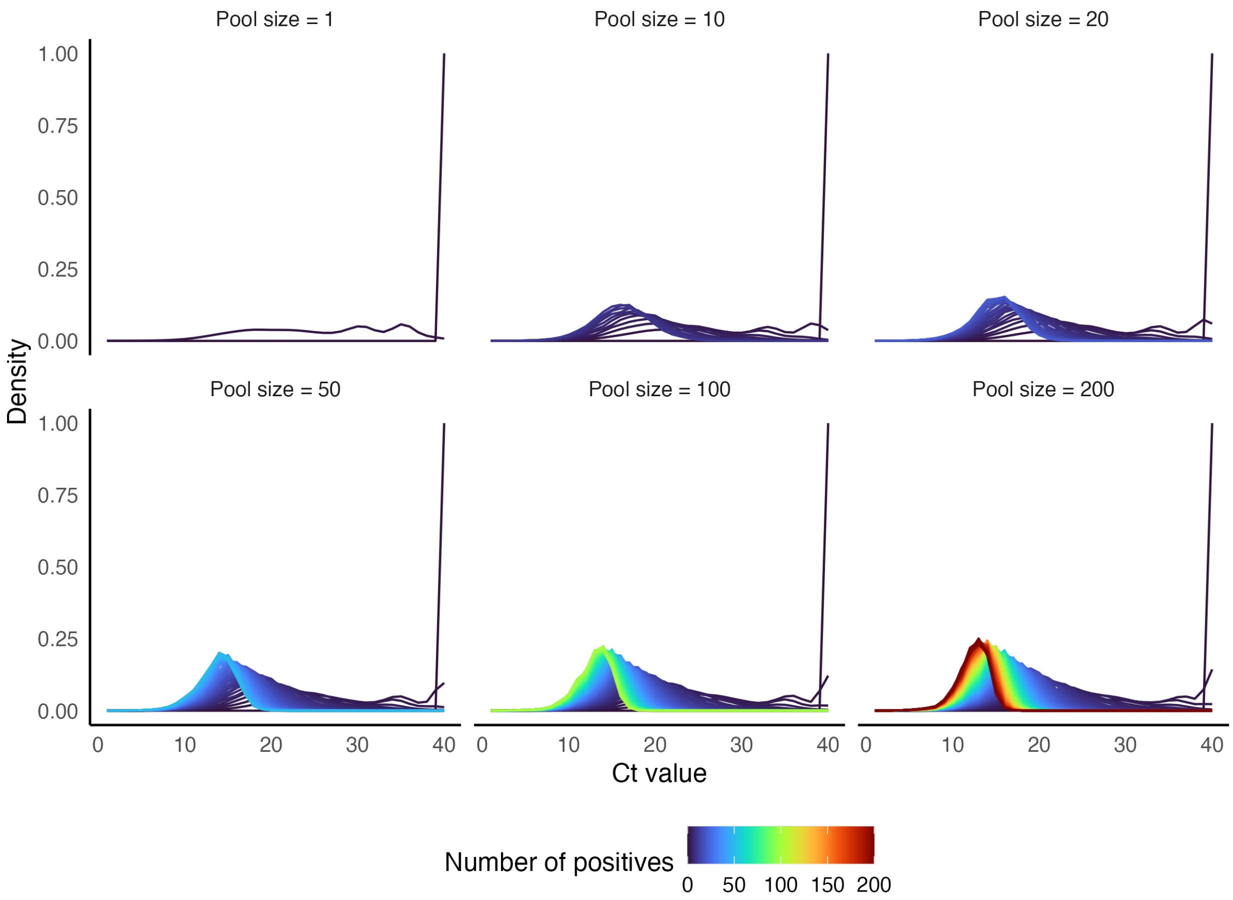

Figure S17: Distributions of estimated $Pr(y_j|k_j, n_j)$ through KDE approximations. Each line represents the number of positive samples ($k_j$). Each panel represents the pool size, and $n_j$ depends on the pool size.

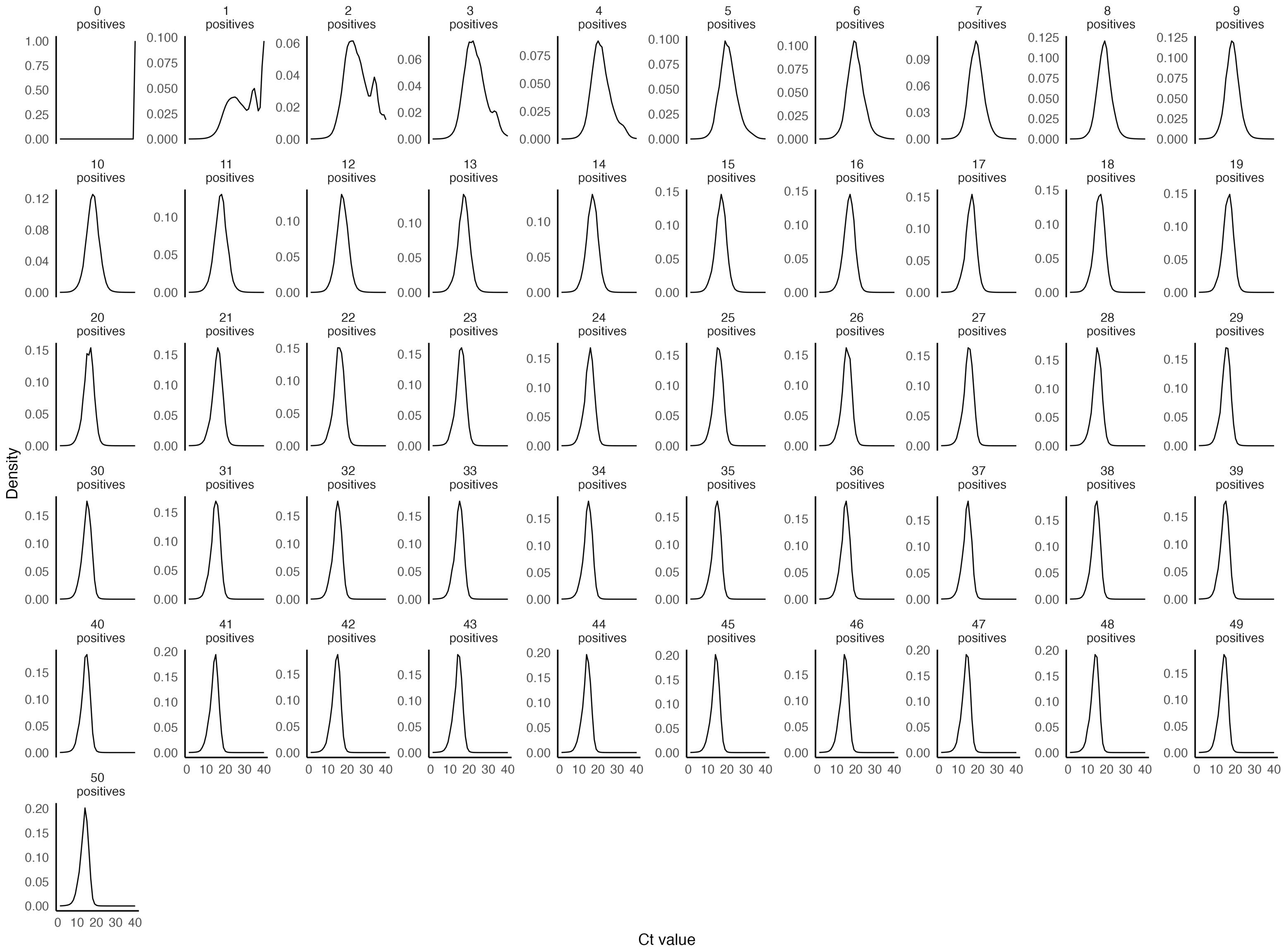

Figure S18: Distributions of estimated $Pr(y_j|k_j, n_j)$ through KDE approximations when the pool size is 50. Each panel represents the number of positive samples ($k_j$). $n_j$ is assumed to be 50.

##### S8.1 Prevalence estimation using other simulated data

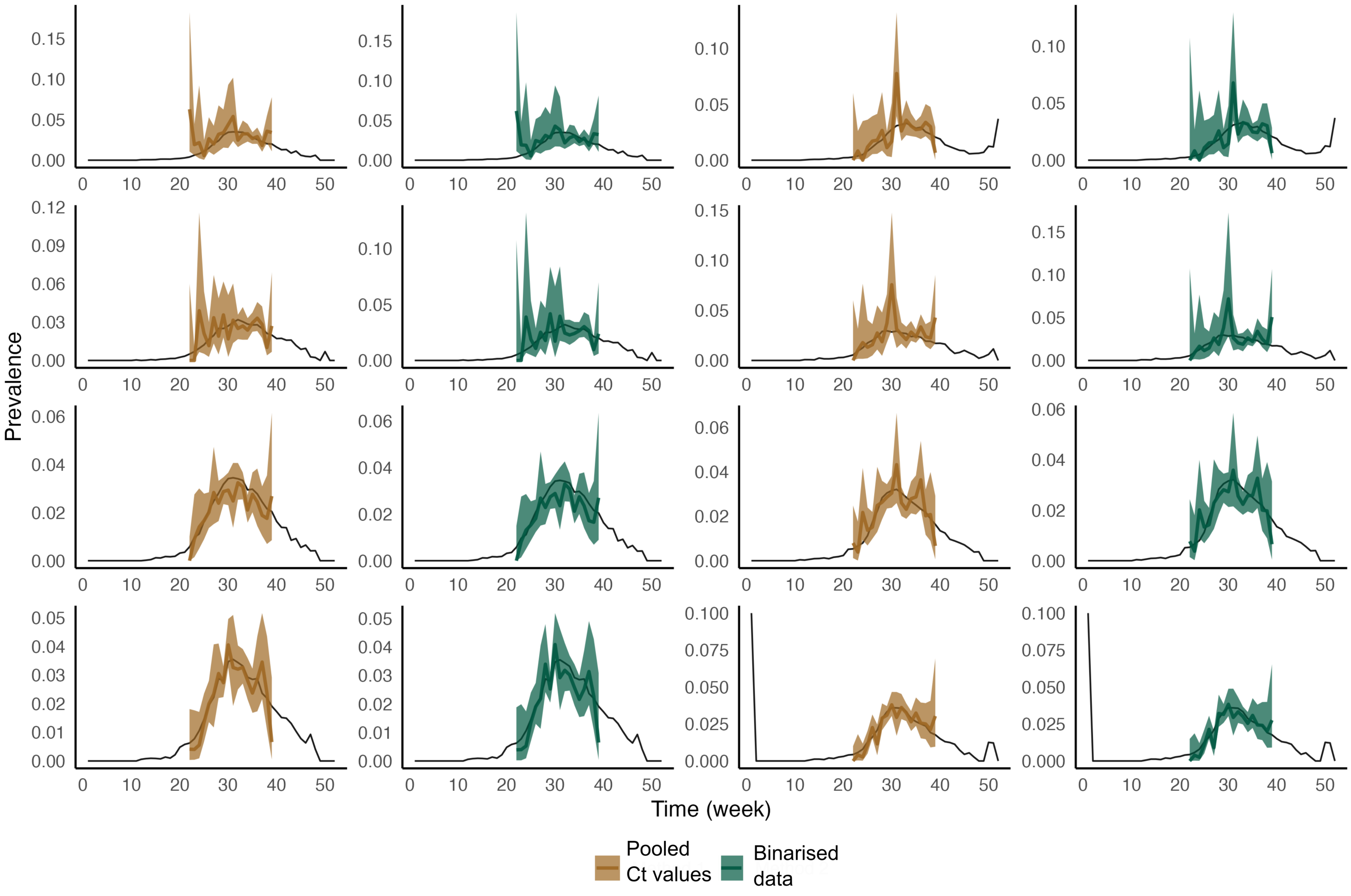

Figure S19: Estimated prevalence using 8 simulations. Beige lines and ribbons illustrate the estimated prevalence and the corresponding 95% confidence intervals using pooled Ct values. The green lines and ribbons illustrate the estimated prevalence and the corresponding 95% confidence intervals using the binarised data.

#### S9 Potential effects of pool size and number of pools on the accuracy of prevalence estimation

In this experiment, we tested the effects of pool size and the number of pools when the true prevalence was in the range $(0.001, 0.8)$. We changed pool sizes from 1, 10, 20, 50, 100 and 200, and the number of pools from 1, 10, 20, 50, 100, and 500. Under each of these combinations, we sampled pools from the Ct values generated from our agent-based model. Then we repeated the pooling and calculation of the pooled Ct values step as described in Section S4.1. The Ct values were binarised based on whether they were equal to 40 (PCR negative) or not (PCR positive).

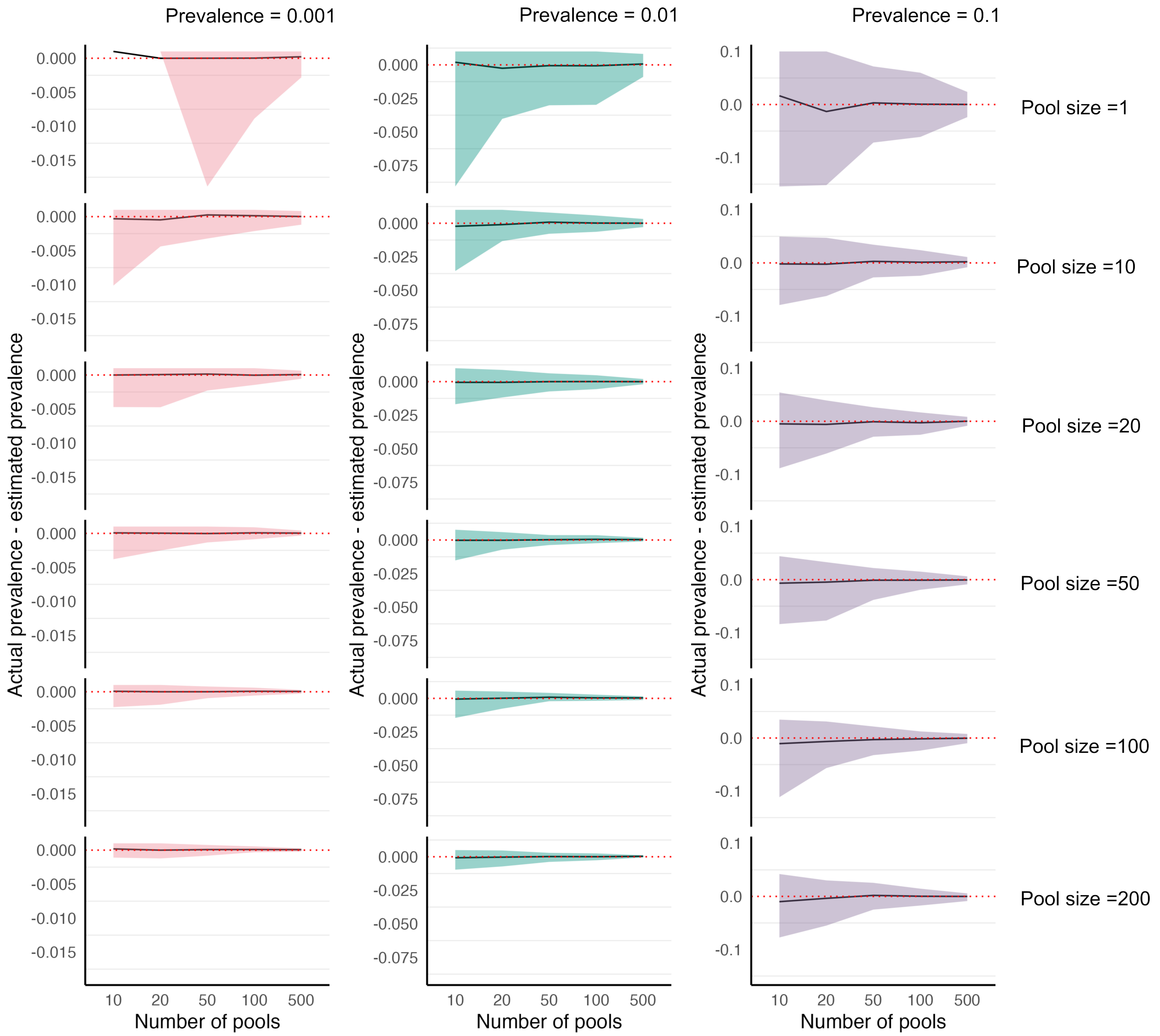

Figure S20: Using Pooled Ct values, the difference between actual prevalence and the estimated prevalence under different prevalence values, pool sizes and number of pools. Each combination was repeated 100 times. All the calculations are done using the MLE estimates for prevalence. Black lines represent the mean of the estimates. The ribbons are 95% quantile intervals based on the MLE estimates. If the MLE estimate is the same across the 100 repeated calculations (see top corner panel when the prevalence =0.001, the quantiles were not calculated.)

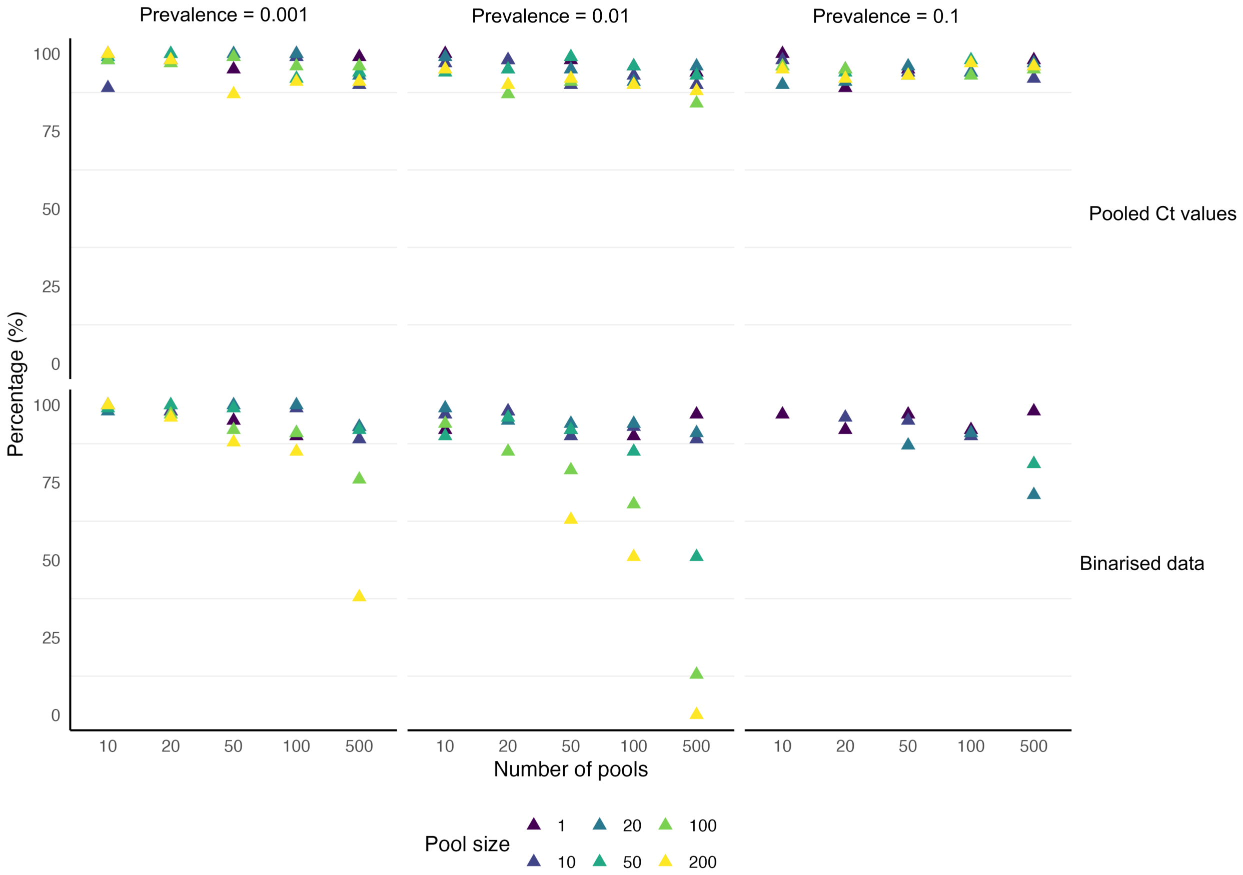

Figure S21: The percentages of 95% confidence intervals that include the true prevalence. 95% confidence intervals were calculated using the same combinations of number of pools, prevalence values, and pool sizes as in Figure S20. Each combination was repeated 100 times, and under each of these measures, each 95% confidence interval was checked to see if the interval covered the true prevalence.

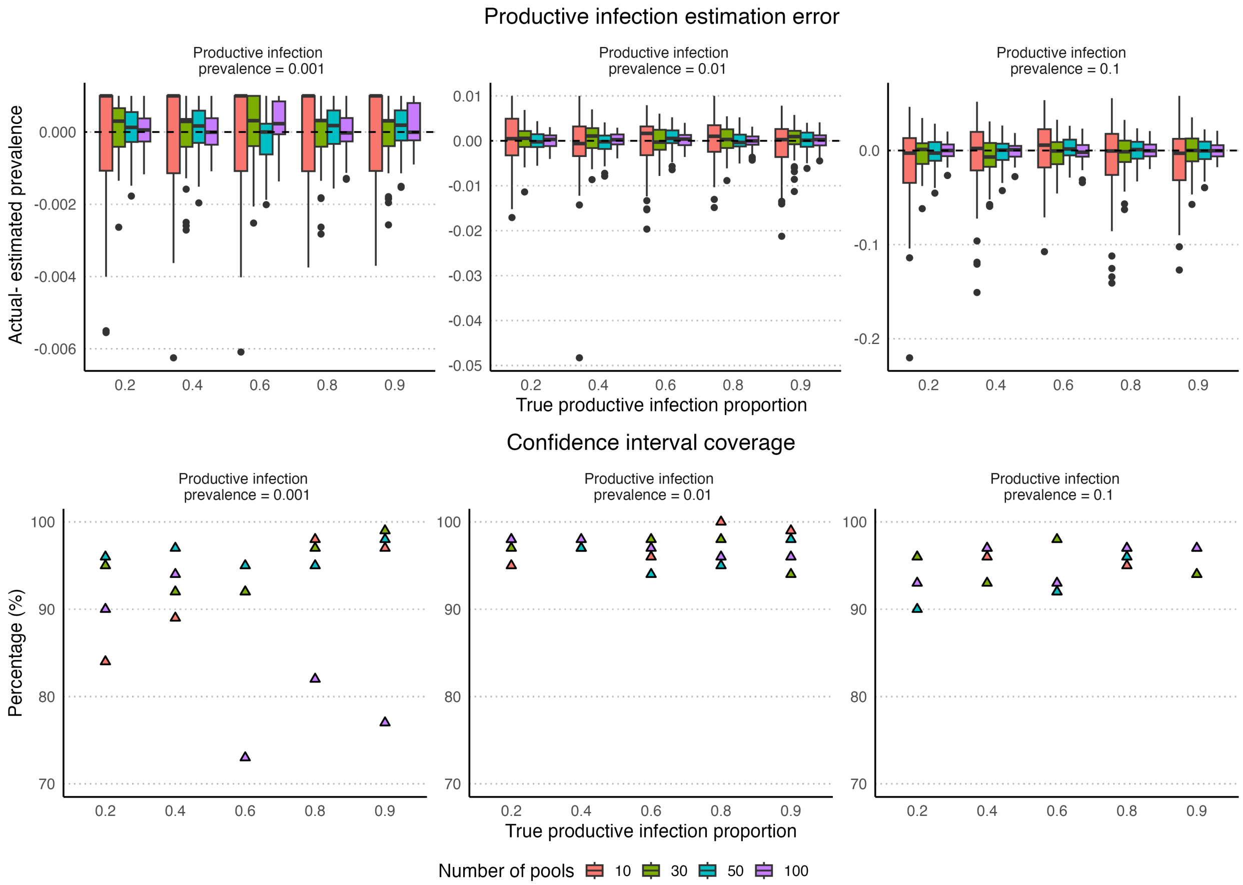

Figure S22: **Top panel: Productive infection estimation error.** Box plots of the difference between actual prevalence and the estimated productive infection prevalence using Pooled Ct values under different prevalence values, number of pools and pool size of 50. Each combination was repeated 100 times. All the calculations are done using the MLE estimates for prevalence.

**Bottom panel: Confidence interval coverage.** The percentages of 95% confidence intervals that include the true productive infection prevalence. 95% confidence intervals were calculated using the same combinations of number of pools, prevalence values, and pool sizes as in top panel. Each combination was repeated 100 times, and under each of these measures, each 95% confidence interval was checked to see if the interval covered the true prevalence.

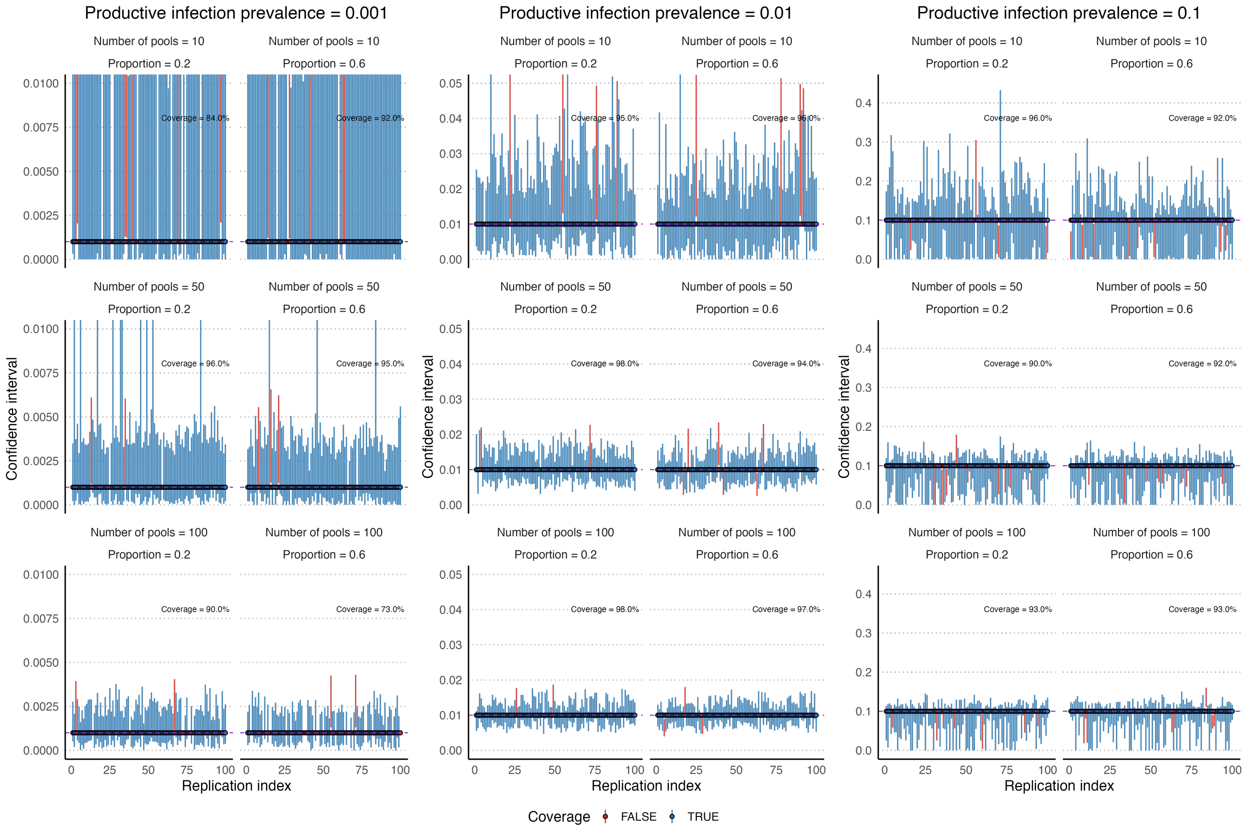

Figure S23: The percentages of 95% confidence intervals that include the true productive infection prevalence. 95% confidence intervals were calculated using the same combinations as in Figure S22. Each combination was repeated 100 times, and under each of these measures, each 95% confidence interval was checked to see if the interval covered the true prevalence. If the interval included the true productive infection prevalence, it is denoted in blue, otherwise in red. Note: Some confidence intervals are narrow (e.g.: see coverage under 100 pools, productive infection proportion 0.6 and production infection prevalence 0.001). Therefore, the confidence intervals may not be visible.

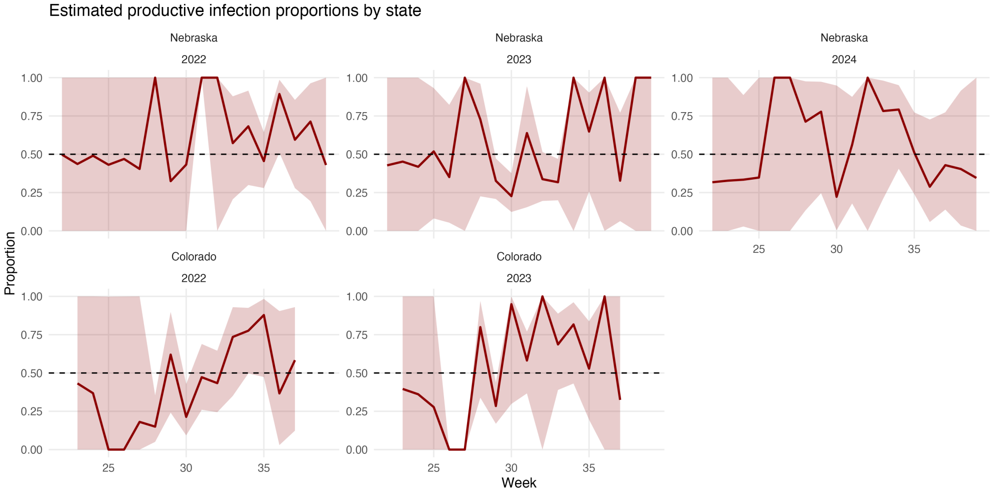

Figure S24: Estimated productive infection proportion from Nebraska and Colorado data based on the results in Figure 7 in the manuscript.

Table S2: Summary statistics for the Ct values from Nebraska from 2022 to 2024. The 95% confidence intervals from both estimation methods partially overlap in all weeks across all years.

| **Year** | **Disease week** | **Number of pools** | **Mean Ct (including negative pools)** | **Minimum Ct** | **Number of positives** | **Mean Ct (with positive pools)** | **Average pool size** | **Estimated prevalence: binary data (95% CI)** | **Estimated prevalence: pooled Ct values (95% CI)** | **Estimated productive infection prevalence: pooled Ct values (95% CI)** |
|---|---|---|---|---|---|---|---|---|---|---|
| 2022 | 22 | 17 | 40.00 | 40.00 | 0 | | 2 | 0 (0, 0.092) | 0 (0, 0.05) | 0 (0, 1) |
| 2022 | 23 | 33 | 40.00 | 40.00 | 0 | | 6 | 0 (0, 0.018) | 0 (0, 0.01) | 0 (0, 1) |
| 2022 | 24 | 25 | 40.00 | 40.00 | 0 | | 3 | 0 (0, 0.045) | 0 (0, 0.025) | 0 (0, 1) |
| 2022 | 25 | 44 | 40.00 | 40.00 | 0 | | 6 | 0 (0, 0.014) | 0 (0, 0.01) | 0 (0, 1) |
| 2022 | 26 | 36 | 40.00 | 40.00 | 0 | | 4 | 0 (0, 0.023) | 0 (0, 0.01) | 0 (0, 1) |
| 2022 | 27 | 36 | 40.00 | 40.00 | 0 | | 7 | 0 (0, 0.014) | 0 (0, 0.01) | 0 (0, 1) |
| 2022 | 28 | 41 | 39.70 | 27.78 | 1 | 27.78 | 6 | 0.004 (0, 0.019) | 0.004 (0, 0.019) | 0.004 (0, 0.021) |
| 2022 | 29 | 45 | 40.00 | 40.00 | 0 | | 11 | 0 (0, 0.008) | 0 (0, 0.005) | 0 (0, 1) |
| 2022 | 30 | 28 | 40.00 | 40.00 | 0 | | 6 | 0 (0, 0.022) | 0 (0, 0.01) | 0 (0, 1) |
| 2022 | 31 | 32 | 39.16 | 21.62 | 2 | 26.52 | 12 | 0.005 (0.001, 0.016) | 0.006 (0.001, 0.018) | 0.005 (0.001, 0.02) |
| 2022 | 32 | 70 | 39.69 | 27.26 | 2 | 29.04 | 22 | 0.001 (0, 0.004) | 0.001 (0, 0.005) | 0.001 (0, 0.004) |
| 2022 | 33 | 60 | 38.50 | 23.27 | 8 | 28.74 | 21 | 0.007 (0.003, 0.014) | 0.008 (0.004, 0.015) | 0.005 (0.002, 0.012) |
| 2022 | 34 | 79 | 38.33 | 20.84 | 10 | 26.79 | 18 | 0.008 (0.004, 0.014) | 0.009 (0.004, 0.015) | 0.006 (0.003, 0.012) |
| 2022 | 35 | 107 | 37.21 | 22.01 | 29 | 29.72 | 29 | 0.012 (0.008, 0.016) | 0.012 (0.008, 0.017) | 0.007 (0.004, 0.01) |
| 2022 | 36 | 92 | 37.05 | 21.20 | 19 | 25.74 | 20 | 0.013 (0.008, 0.02) | 0.015 (0.009, 0.022) | 0.012 (0.007, 0.019) |
| 2022 | 37 | 74 | 37.96 | 19.40 | 11 | 26.26 | 18 | 0.01 (0.005, 0.017) | 0.011 (0.006, 0.019) | 0.007 (0.004, 0.014) |
| 2022 | 38 | 66 | 38.88 | 19.13 | 5 | 25.21 | 13 | 0.006 (0.002, 0.013) | 0.007 (0.002, 0.015) | 0.005 (0.002, 0.013) |
| 2022 | 39 | 41 | 40.00 | 40.00 | 0 | | 5 | 0 (0, 0.018) | 0 (0, 0.01) | 0 (0, 1) |
| 2023 | 22 | 53 | 40.00 | 40.00 | 0 | | 6 | 0 (0, 0.013) | 0 (0, 0.005) | 0 (0, 1) |
| 2023 | 23 | 35 | 40.00 | 40.00 | 0 | | 6 | 0 (0, 0.017) | 0 (0, 0.01) | 0 (0, 1) |
| 2023 | 24 | 53 | 40.00 | 40.00 | 0 | | 6 | 0 (0, 0.011) | 0 (0, 0.005) | 0 (0, 1) |
| 2023 | 25 | 58 | 39.40 | 20.00 | 3 | 28.37 | 13 | 0.004 (0.001, 0.011) | 0.005 (0.001, 0.012) | 0.003 (0.001, 0.011) |
| 2023 | 26 | 105 | 39.60 | 25.20 | 4 | 29.48 | 27 | 0.001 (0, 0.003) | 0.002 (0, 0.004) | 0.001 (0, 0.003) |
| 2023 | 27 | 44 | 39.71 | 27.10 | 1 | 27.10 | 15 | 0.002 (0, 0.007) | 0.002 (0, 0.008) | 0.002 (0, 0.008) |
| 2023 | 28 | 92 | 38.63 | 22.00 | 10 | 27.39 | 22 | 0.006 (0.003, 0.01) | 0.006 (0.003, 0.011) | 0.005 (0.002, 0.009) |
| 2023 | 29 | 159 | 37.42 | 20.00 | 42 | 30.22 | 36 | 0.009 (0.006, 0.012) | 0.009 (0.007, 0.012) | 0.004 (0.003, 0.006) |
| 2023 | 30 | 135 | 38.05 | 19.50 | 32 | 31.77 | 32 | 0.009 (0.006, 0.013) | 0.009 (0.007, 0.013) | 0.003 (0.002, 0.006) |
| 2023 | 31 | 51 | 39.01 | 21.90 | 4 | 27.38 | 15 | 0.006 (0.002, 0.014) | 0.007 (0.002, 0.015) | 0.004 (0.001, 0.014) |
| 2023 | 32 | 128 | 37.65 | 17.80 | 30 | 29.97 | 26 | 0.011 (0.008, 0.016) | 0.012 (0.008, 0.016) | 0.005 (0.003, 0.008) |

| | | | | | | | | | | |
|---|---|---|---|---|---|---|---|---|---|---|
| 2023 | 33 | 145 | 37.28 | 19.00 | 41 | 30.36 | 39 | 0.009 (0.006, 0.012) | 0.009 (0.007, 0.013) | 0.004 (0.003, 0.006) |
| 2023 | 34 | 129 | 38.14 | 20.70 | 17 | 25.86 | 32 | 0.005 (0.003, 0.007) | 0.005 (0.003, 0.008) | 0.004 (0, 0.007) |
| 2023 | 35 | 85 | 38.28 | 16.80 | 9 | 23.72 | 27 | 0.004 (0.002, 0.007) | 0.005 (0.002, 0.008) | 0.003 (0.002, 0.007) |
| 2023 | 36 | 60 | 38.16 | 19.84 | 7 | 24.24 | 9 | 0.013 (0.006, 0.026) | 0.014 (0.006, 0.028) | 0.013 (0, 0.024) |
| 2023 | 37 | 54 | 39.08 | 19.20 | 5 | 30.08 | 17 | 0.006 (0.002, 0.012) | 0.006 (0.002, 0.013) | 0.002 (0.001, 0.009) |
| 2023 | 38 | 56 | 39.26 | 19.14 | 2 | 19.39 | 8 | 0.004 (0.001, 0.014) | 0.005 (0.001, 0.015) | 0.004 (0, 0.014) |
| 2023 | 39 | 38 | 39.59 | 24.30 | 1 | 24.30 | 5 | 0.005 (0, 0.022) | 0.005 (0, 0.023) | 0.005 (0, 0.028) |
| 2024 | 22 | 57 | 40.00 | 40.00 | 0 | | 15 | 0 (0, 0.005) | 0 (0, 0.005) | 0 (0, 1) |
| 2024 | 23 | 53 | 40.00 | 40.00 | 0 | | 13 | 0 (0, 0.005) | 0 (0, 0.005) | 0 (0, 1) |
| 2024 | 24 | 84 | 39.71 | 20.40 | 2 | 27.85 | 24 | 0.001 (0, 0.003) | 0.001 (0, 0.003) | 0 (0, 0.003) |
| 2024 | 25 | 51 | 40.00 | 40.00 | 0 | | 18 | 0 (0, 0.004) | 0 (0, 0.005) | 0 (0, 1) |
| 2024 | 26 | 73 | 39.80 | 25.28 | 1 | 25.28 | 13 | 0.001 (0, 0.005) | 0.001 (0, 0.005) | 0.001 (0, 0.005) |
| 2024 | 27 | 50 | 39.52 | 27.30 | 2 | 28.00 | 11 | 0.004 (0.001, 0.012) | 0.004 (0.001, 0.013) | 0.004 (0, 0.012) |
| 2024 | 28 | 111 | 39.45 | 21.30 | 5 | 27.80 | 23 | 0.002 (0.001, 0.004) | 0.002 (0.001, 0.005) | 0.002 (0.001, 0.004) |
| 2024 | 29 | 89 | 38.81 | 21.00 | 8 | 26.76 | 30 | 0.003 (0.001, 0.006) | 0.004 (0.002, 0.007) | 0.003 (0.001, 0.006) |
| 2024 | 30 | 77 | 39.80 | 30.90 | 2 | 32.40 | 21 | 0.001 (0, 0.004) | 0.001 (0, 0.004) | 0 (0, 0.008) |
| 2024 | 31 | 60 | 38.74 | 23.20 | 7 | 29.17 | 20 | 0.007 (0.003, 0.013) | 0.007 (0.003, 0.014) | 0.004 (0.002, 0.011) |
| 2024 | 32 | 64 | 38.71 | 21.90 | 5 | 23.50 | 16 | 0.005 (0.002, 0.012) | 0.006 (0.002, 0.013) | 0.005 (0, 0.011) |
| 2024 | 33 | 75 | 38.31 | 19.80 | 9 | 25.94 | 25 | 0.005 (0.002, 0.009) | 0.006 (0.003, 0.01) | 0.004 (0.002, 0.009) |
| 2024 | 34 | 105 | 37.34 | 18.70 | 18 | 24.47 | 29 | 0.007 (0.004, 0.01) | 0.008 (0.005, 0.012) | 0.006 (0.004, 0.01) |
| 2024 | 35 | 108 | 38.06 | 23.10 | 20 | 29.52 | 29 | 0.008 (0.005, 0.012) | 0.008 (0.005, 0.012) | 0.005 (0.003, 0.009) |
| 2024 | 36 | 80 | 39.34 | 20.00 | 5 | 29.48 | 19 | 0.003 (0.001, 0.007) | 0.004 (0.001, 0.008) | 0.001 (0, 0.006) |
| 2024 | 37 | 63 | 38.73 | 20.90 | 7 | 28.60 | 17 | 0.007 (0.003, 0.014) | 0.008 (0.003, 0.015) | 0.004 (0.002, 0.011) |
| 2024 | 38 | 51 | 39.59 | 21.70 | 2 | 29.55 | 9 | 0.004 (0.001, 0.014) | 0.005 (0.001, 0.014) | 0.002 (0, 0.016) |
| 2024 | 39 | 52 | 40.00 | 40.00 | 1 | | 10 | 0 (0, 0.008) | 0 (0, 0.005) | 0 (0, 1) |

Table S3: Summary statistics for the Ct values from Colorado from 2022-2023. The 95% confidence intervals from both estimation methods partially overlap in all weeks across all years.

| Year | Disease week | Number of pools | Mean Ct (including negative pools) | Minimum Ct | Number of positives | Mean Ct (with positive pools) | Average pool size | Estimated prevalence: binary data (95% CI) | Estimated prevalence: pooled Ct values (95% CI) | Estimated productive infection prevalence: pooled Ct values (95% CI) |
|---|---|---|---|---|---|---|---|---|---|---|
| 2022 | 23 | 41 | 40.00 | 40.00 | 0 | | 7 | 0 (0, 0.014) | 0 (0, 0.005) | 0 (0, 1) |
| 2022 | 24 | 62 | 40.00 | 40.00 | 0 | | 8 | 0 (0, 0.007) | 0 (0, 0.005) | 0 (0, 1) |
| 2022 | 25 | 60 | 39.92 | 37.57 | 2 | 37.65 | 9 | 0.004 (0.001, 0.012) | 0.004 (0.001, 0.013) | 0 (0, 0.998) |
| 2022 | 26 | 85 | 39.83 | 32.49 | 2 | 32.73 | 14 | 0.002 (0, 0.005) | 0.002 (0, 0.005) | 0 (0, 0.999) |
| 2022 | 27 | 117 | 40.00 | 40.00 | 0 | | 24 | 0 (0, 0.001) | 0 (0, 0.005) | 0 (0, 1) |
| 2022 | 28 | 164 | 39.22 | 19.45 | 15 | 31.47 | 32 | 0.003 (0.002, 0.005) | 0.003 (0.002, 0.005) | 0.001 (0, 0.002) |
| 2022 | 29 | 176 | 39.27 | 19.76 | 9 | 25.66 | 33 | 0.002 (0.001, 0.003) | 0.002 (0.001, 0.003) | 0.001 (0.001, 0.003) |
| 2022 | 30 | 139 | 38.84 | 17.31 | 18 | 31.04 | 26 | 0.005 (0.003, 0.008) | 0.006 (0.003, 0.009) | 0.002 (0.001, 0.004) |
| 2022 | 31 | 130 | 38.10 | 19.25 | 19 | 27.01 | 26 | 0.006 (0.004, 0.009) | 0.007 (0.004, 0.01) | 0.004 (0.002, 0.007) |
| 2022 | 32 | 144 | 37.70 | 8.41 | 22 | 24.97 | 31 | 0.006 (0.004, 0.008) | 0.006 (0.004, 0.009) | 0.003 (0.002, 0.006) |
| 2022 | 33 | 147 | 38.71 | 19.10 | 13 | 25.40 | 28 | 0.003 (0.002, 0.006) | 0.004 (0.002, 0.006) | 0.003 (0.002, 0.005) |
| 2022 | 34 | 170 | 37.63 | 18.34 | 25 | 23.88 | 30 | 0.005 (0.003, 0.008) | 0.006 (0.004, 0.009) | 0.005 (0.003, 0.007) |
| 2022 | 35 | 113 | 37.50 | 17.46 | 18 | 24.31 | 17 | 0.011 (0.007, 0.017) | 0.012 (0.007, 0.018) | 0.01 (0.006, 0.016) |
| 2022 | 36 | 92 | 39.77 | 24.09 | 2 | 29.46 | 13 | 0.002 (0, 0.005) | 0.002 (0, 0.005) | 0.001 (0, 0.006) |
| 2022 | 37 | 94 | 39.51 | 23.66 | 4 | 28.58 | 19 | 0.002 (0.001, 0.006) | 0.003 (0.001, 0.006) | 0.002 (0.001, 0.005) |
| 2023 | 23 | 69 | 40.00 | 40.00 | 0 | | 6 | 0 (0, 0.009) | 0 (0, 0.005) | 0 (0, 1) |
| 2023 | 24 | 78 | 40.00 | 40.00 | 0 | | 7 | 0 (0, 0.007) | 0 (0, 0.005) | 0 (0, 1) |
| 2023 | 25 | 107 | 40.00 | 40.00 | 0 | | 17 | 0 (0, 0.002) | 0 (0, 0.005) | 0 (0, 1) |
| 2023 | 26 | 231 | 39.96 | 30.12 | 1 | 30.12 | 39 | 0 (0, 0) | 0 (0, 0.001) | 0 (0, 0) |
| 2023 | 27 | 290 | 39.63 | 19.28 | 8 | 26.65 | 41 | 0.001 (0, 0.001) | 0.001 (0, 0.001) | 0 (0, 0) |
| 2023 | 28 | 322 | 39.26 | 17.15 | 17 | 25.97 | 42 | 0.001 (0.001, 0.002) | 0.001 (0.001, 0.002) | 0.001 (0.001, 0.002) |
| 2023 | 29 | 403 | 39.02 | 14.13 | 37 | 29.28 | 45 | 0.002 (0.002, 0.003) | 0.002 (0.002, 0.003) | 0.001 (0.001, 0.002) |
| 2023 | 30 | 206 | 37.97 | 18.85 | 26 | 23.93 | 36 | 0.004 (0.003, 0.005) | 0.004 (0.003, 0.006) | 0.004 (0.003, 0.005) |
| 2023 | 31 | 142 | 37.05 | 17.70 | 28 | 25.04 | 28 | 0.008 (0.006, 0.012) | 0.009 (0.006, 0.013) | 0.006 (0.004, 0.009) |
| 2023 | 32 | 118 | 36.93 | 16.59 | 21 | 22.73 | 22 | 0.01 (0.006, 0.015) | 0.011 (0.007, 0.017) | 0.01 (0, 0.014) |
| 2023 | 33 | 107 | 37.39 | 18.78 | 20 | 26.01 | 20 | 0.011 (0.007, 0.017) | 0.013 (0.008, 0.019) | 0.009 (0.006, 0.015) |
| 2023 | 34 | 116 | 37.77 | 19.34 | 16 | 23.80 | 19 | 0.008 (0.005, 0.013) | 0.009 (0.005, 0.014) | 0.007 (0.004, 0.012) |
| 2023 | 35 | 96 | 39.05 | 19.20 | 8 | 28.59 | 10 | 0.009 (0.004, 0.017) | 0.01 (0.004, 0.018) | 0.006 (0.002, 0.014) |
| 2023 | 36 | 58 | 39.60 | 16.56 | 1 | 16.56 | 8 | 0.002 (0, 0.009) | 0.002 (0, 0.009) | 0.002 (0, 0.012) |
| 2023 | 37 | 44 | 40.00 | 40.00 | 0 | | 10 | 0 (0, 0.008) | 0 (0, 0.005) | 0 (0, 1) |

#### S10 Availability of codes and related data

Please refer to the link https://github.com/PunyaAlahakoon/west_nile_virus_abm.git.
